# Paternally inherited noncoding structural variants contribute to autism

**DOI:** 10.1101/102327

**Authors:** William M. Brandler, Danny Antaki, Madhusudan Gujral, Morgan L. Kleiber, Michelle S. Maile, Oanh Hong, Timothy R. Chapman, Shirley Tan, Prateek Tandon, Timothy Pang, Shih C. Tang, Keith K. Vaux, Yan Yang, Eoghan Harrington, Sissel Juul, Daniel J. Turner, Stephen F. Kingsmore, Joseph G. Gleeson, Boyko Kakaradov, Amalio Telenti, J Craig Venter, Roser Corominas, Bru Cormand, Isabel Rueda, Karen S. Messer, Caroline M. Nievergelt, Maria J. Arranz, Eric Courchesne, Karen Pierce, Alysson R. Muotri, Lilia M. Iakoucheva, Amaia Hervas, Christina Corsello, Jonathan Sebat

**Affiliations:** Beyster Center for Genomics of Psychiatric Diseases, University of California San Diego, La Jolla, CA, 92093 USA.; Department of Psychiatry, University of California San Diego, La Jolla, CA, 92093 USA.; Department of Cellular and Molecular Medicine and Pediatrics, University of California San Diego, La Jolla, CA, 92093 USA.; Biomedical Sciences Graduate Program, University of California San Diego, La Jolla, CA, 92093 USA.; Rady Children’s Hospital, La Jolla, CA, 92123 USA.; Department of Medicine, University of California San Diego, La Jolla, CA, 92093 USA.; Oxford Nanopore Technologies Inc., New York, NY, 10013 USA.; Oxford Nanopore Technologies Inc., Oxford, UK.; Rady Children Institute for Genomic Medicine, Rady Children Hospital, San Diego, CA, 92123 USA.; Howard Hughes Medical Institute, Rady Children Institute of Genomic Medicine, Department of Neurosciences, University of California San Diego, San Diego, CA, 92093 USA.; Human Longevity Inc., San Diego, CA, 92121 USA.; J. Craig Venter Institute, La Jolla, CA, 92037 USA.; Genetics Research Unit, Universitat Pompeu Fabra, Hospital del Mar Research Institute (IMIM), Barcelona, Spain.; Centro de Investigación Biomédica en Red de Enfermedades Raras (CIBERER), Madrid, Spain.; Institut de Biomedicina de la Universitat de Barcelona (IBUB), Catalonia, Spain.; Departament de Genètica, Facultat de Biologia, Universitat de Barcelona, Catalonia, Spain.; Institut de Recerca Sant Joan de Dèu (IR-SJD), Espluges, Catalonia, Spain.; Department of Psychiatry, Hospital Sant Joan de Deu, Barcelona, Spain.; Division of Biostatistics and Bioinformatics, Department of Family Medicine and Public Health, University of California San Diego, San Diego, CA, 92093 USA.; Research Laboratory Unit, Fundacio Docencia I Recerca Mútua Terrassa, Barcelona, Spain.; Department of Neuroscience, University of California San Diego, La Jolla, CA, 92093 USA.; Child and Adolescent Mental Health Unit, Hospital Universitari Mútua de Terrassa, Barcelona, Spain.

**Keywords:** genomics, whole genome sequencing, autism, noncoding, mosaics, structural variation

## Abstract

The genetic architecture of autism spectrum disorder (ASD) is known to consist of contributions from gene-disrupting de novo mutations and common variants of modest effect. We hypothesize that the unexplained heritability of ASD also includes rare inherited variants with intermediate effects. We investigated the genome-wide distribution and functional impact of structural variants (SVs) through whole genome analysis (*≥*30X coverage) of 3,169 subjects from 829 families affected by ASD. Genes that are intolerant to inactivating variants in the exome aggregation consortium (ExAC) were depleted for SVs in parents, specifically within fetal-brain promoters, UTRs and exons. Rare paternally-inherited SVs that disrupt promoters or UTRs were over-transmitted to probands (*P* = 0.0013) and not to their typically-developing siblings. Recurrent functional noncoding deletions implicate the gene *LEO1* in ASD. Protein-coding SVs were also associated with ASD (*P* = 0.0025). Our results establish that rare inherited SVs predispose children to ASD, with differing contributions from each parent.

## Introduction

Autism Spectrum Disorders (ASDs) have a complex etiology with a major contribution from genetic factors. Microarray and exome sequencing studies over the past decade have demonstrated that de novo gene-disrupting or protein-altering variants contribute in approximately 25% of cases [1, 2, 3, 4, 5, 6, 7, 8, 9], and causality has been demonstrated for many genes [10]. In addition, common variants of modest effect contribute to risk for ASD [11]. Thus, the genetic architecture of ASD consists of a wide spectrum of risk alleles. At one extreme are the dominant-acting variants that carry high risk and are rarely carried by asymptomatic parents. At the opposite extreme are many common ‘polygenes’ which individually exert subtle influences on risk.

Much of the allelic spectrum of ASD genetics however has been unexplored, namely rare inherited coding or noncoding variants with intermediate effects [12]. Recent studies have developed our understanding how much of the genome is regulatory through analyses of evolutionarily conservation and identification of biochemically active noncoding genetic elements [13, 14]. However, functional noncoding variants are not easily distinguishable from the vast background of neutral variation in the general population. Initial applications of whole genome sequencing (WGS) in ASD therefore have been underpowered to detect any association of rare noncoding point mutations with ASD [15, 16, 17, 18].

Structural variants (SVs), such as deletions, duplications, insertions and inversions [19], are more likely to impact gene regulation because of their potential to disrupt, duplicate, and shuffle functional elements in the genome. SVs could therefore provide a foothold for expanding our knowledge of the genetic architecture of ASD beyond what is detectable through exome sequencing or GWAS. Recent WGS efforts led by the 1000 Genomes consortium and our group have revealed thousands of rare, inherited SVs in each genome that were previously undetectable with microarray or exome sequencing technologies [19, 20].

We hypothesize rare, inherited SVs that disrupt functional elements of variant-intolerant genes critical for neurodevelopment will be enriched for variants that contribute to autism spectrum disorder. In order to assess this we have created a pipeline for accurate detection and genotyping of SVs in high coverage WGS data and investigated genetic association across multiple classes of variants (inherited, de novo, coding and noncoding) in two independent cohorts comprising 3,169 individuals from 829 families affected by ASD. We find that paternally inherited noncoding CNVs that disrupt promoters or UTRs of variant-intolerant genes are preferentially transmitted to affected offspring and not to their unaffected siblings, replicating this finding in both cohorts, and implicating the novel gene *LEO1* in ASD.

## Results

### Genome-wide detection and genotyping of SV in ASD families

We investigated SVs genome wide by high coverage whole genome sequencing (mean coverage = 42.6) of 3,169 individuals from two cohorts: (1) the REACH cohort consisted of 311 families with 362 affected offspring and 112 sibling controls (*n* = 1,097 genomes) recruited from Hospitals and clinics in San Diego and Barcelona and sequenced in San Diego and (2) The Simons Simplex Collection (SSC) dataset consisted of 518 discordant sibling-pair quad families (*n* = 2,072 genomes) sequenced at New York Genome Center. By design, these two cohorts differ slightly with respect to genetic etiology. The REACH cohort is a representative sample of ASD, and had not been previously analyzed by microarrays or exome sequencing. The ratio of males to females in cases was 4:1 in the REACH cohort. The SSC sample was selected from a larger cohort of 2,644 families [7, 9] after excluding families in which cases or sibling controls carried a large de novo copy number variant (CNV) or truncating point mutation from microarray or exome sequencing. Thus, the SSC cohort was selected to enrich for novel genetic etiology and has a diminished contribution from de novo mutations that are detectable by standard genetic approaches. The SSC sample was disproportionately male (8:1), which was in part due to the removal of de novo mutation carriers that tend to be overrepresented in females (a lower male-female ratio of 2:1) [21]. In total 829 families were sequenced, comprising 880 affected, 630 unaffected individuals, and their parents (**Table S1**).

We developed a pipeline for genome wide analysis of SV that consisted of multiple complementary methods for SV discovery combined with custom software for estimating genotype likelihoods from the combined set of SV calls (Figure S1). To assess the association of inherited SVs with ASD in families, high genotyping accuracy is needed [22]. Thus, a key innovation of the current pipeline was the development and refinement of SV^2^, a support-vector machine (SVM) based software for estimating genotype likelihoods from short read WGS data [23]. Genotype likelihoods serve as our primary metric for SV filtering and assigning of SV genotypes in families. The genotyping accuracy of SV^2^ and the potential for spurious associations to arise from genotyping error was evaluated in this study as part of a companion paper [23].

Briefly, the primary variant calls include biallelic deletions and tandem duplications, inversions, four classes of complex SVs, reciprocal translocations, and four classes of mobile element insertion. An average of 3,746 SVs were detected per individual, the majority of which were deletions (2,428 / individual), *Alu* insertions (920 / individual) and tandem duplications (174 / individual; **Table S2**). Variants were typically private to individual families, being present in only one parent (53.1%), although 48.8% overlapped (≥50% reciprocally) with variants from the 1000 Genomes Phase 3 callset (**Figures S2 and S3**). False discovery rates (FDR) for deletion and duplication calls were estimated from Illumina 2.5M SNP array data (using SVToolkit, see Materials and Methods), which was collected on a subset of samples in our study (*n* = 205). FDR was estimated to be 4.2% for deletions, 9.4% for duplications (Figure S4; **Table S3**), and 6.5% for deletions and duplications within complex SVs. We demonstrate that private deletions and duplications >100bp in size have low FDR and Mendelian-error rates and neutral transmission to offspring (Figure S4). Given that deletions and duplications *>*100 bp comprise the majority of SV calls, can be uniformly genotyped with high accuracy, and their functional impact is more readily interpretable, our subsequent analyses was restricted to this subset of SVs.

### Defining genomic elements depleted for structural variation

The prioritization of rare variants in disease studies is aided by knowledge of the functional elements and genes that are under functional constraint in humans, as illustrated by recent studies that utilize variant frequencies from the exome aggregation consortium (ExAC) to prioritize genes for disease studies [24]. We expect that the genetic effect of rare inherited and de novo variants will be most readily detectable among functional elements that display a demonstrable signature of negative selection for SVs. To this end we sought to define classes of cis-regulatory elements that are depleted in rare SVs in parents in our dataset compared to a random distribution of SVs based on permutations. Noncoding SVs were defined as SVs that did not intersect with any protein coding exons. Considering all genes, we found a depletion of deletions in protein coding exons (odds ratio OR = 0.46; *P* < 0.0001), and variants disrupting 5’UTRs and transcription start sites (TSS OR = 0.77; *P* = 0.0003), and 3’UTRs (OR = 0.87; *P* = 0.007) (**Table S4**), relative to permuted SVs. All other features showed no significant depletion of SV after FDR adjustment for 27 features tested (**Table S4**).

**Figure 1.**
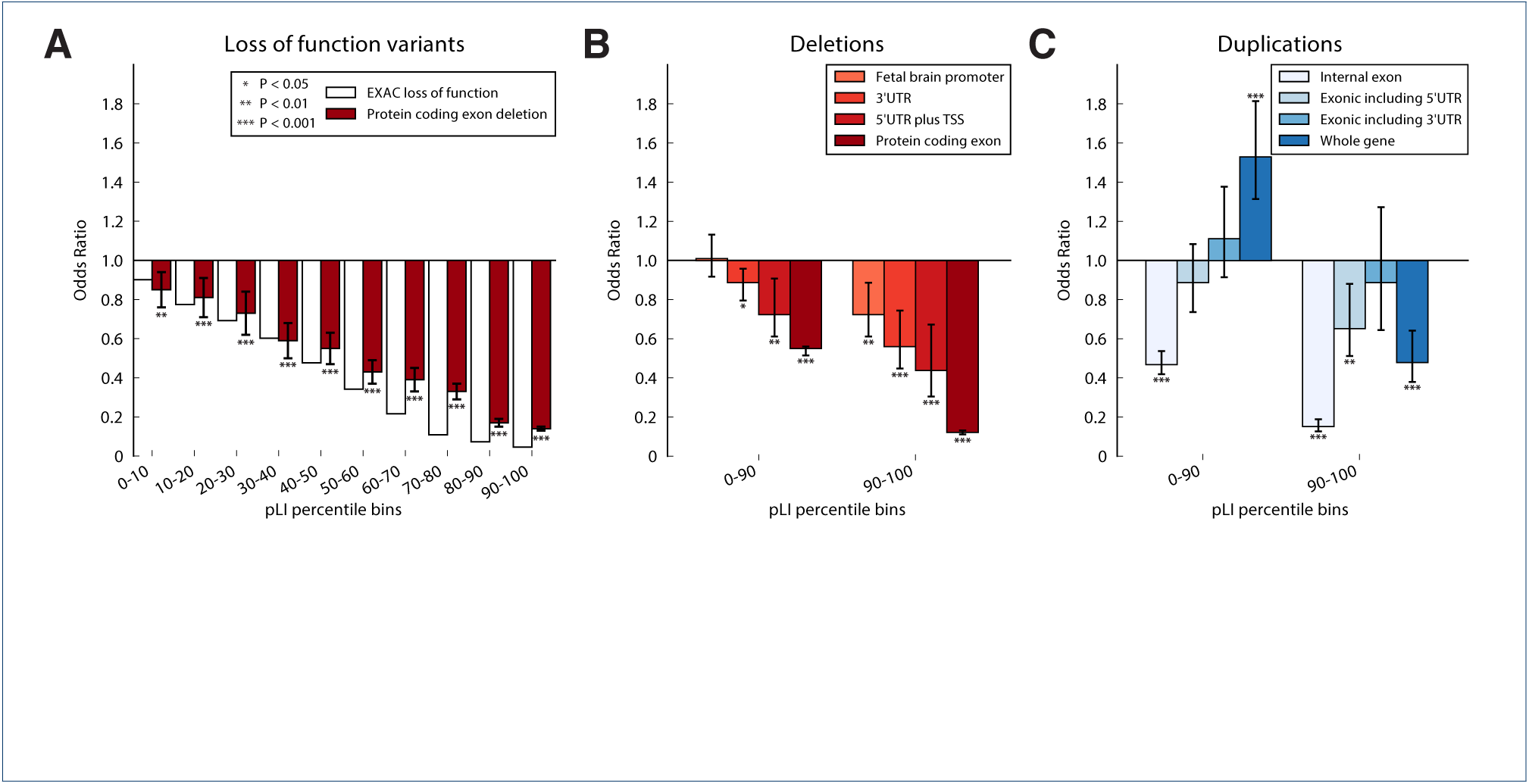
Classes of functional element that are intolerant to structural variation. Bar charts illustrating functional elements that show depletions in structural variation relative to random permutations, stratified on variant-intolerance of genes as estimated by the EXAC consortium (see **Table S4** for a comprehensive set of genomic elements). A) Odds ratios for loss of function mutations from EXAC compared to protein coding deletions from our study, binning genes based on their probability of loss of function intolerance (pLI) scores. B) Coding and noncoding features depleted in deletions. C) Odds ratios highlighting the enrichment/depletion of four different classes of exonic duplication. The number of stars indicates the level of significance; whiskers represent 95% confidence intervals.

The depletion of SVs within functional elements correlated with independent measures of the functional constraint of genes from ExAC (**Table S4**; Figure 1) [24]. ExAC contains a collection of 46,785 exomes from individuals who do not have psychiatric disorders, which has been used to identify genes that are depleted in loss of function mutations [24]. Binning genes by ExAC probability of loss-of-function intolerance (pLI) scores, there was a positive correlation between depletion of exonic deletions and depletion of loss of function point mutations (Figure 1A; Pearson’s r = 0.98). Considering only genes with pLI scores in the 90^th^ percentile or greater, we observed a significant depletion of SVs in exons, UTRs and TSSs. In addition, chromatin marks associated with promoters in fetal brain tissue also showed depletion among these variant-intolerant genes (OR = 0.73; *P* = 0.0011).

We divided exonic duplications into four categories; whole gene duplications, internal exon duplications, exonic duplications that also duplicate the 5’UTR (but not 3’UTR), and exonic duplications that include the 3’UTR (but not 5’UTR; Figure S5). Whole gene duplications were depleted if they duplicated the most variant-intolerant genes (pLI ≥ 90^th^ percentile; OR = 0.49; *P* < 0.0001; Figure 1C) and enriched in genes that are tolerant to inactivating mutations (pLI *<*90^th^ percentile; OR = 1.50; *P* < 0.0001). Internal exon duplications showed depletions similar to that of exonic deletions, consistent with their predicted loss of function effect (Figure 1C). Exonic duplications that encompassed the 5’UTR were also depleted in the most variant-intolerant genes (pLI ≥ 90^th^ percentile OR = 0.68; *P* = 0.007; Figure 1C), but 3’UTR exonic duplications showed no depletion (**Table S5**; Figure 1C). Functional classes of SV that were most strongly depleted in the genome relative to permutations (Fig 1B-C) were subsequently selected for family-based association tests including deletions of fetal brain promoters, UTRs, TSSs and exons, and duplications of UTRs or exons in variant-intolerant genes. The same loci are also highly enriched in known autism genes from exome and CNV studies (OR = 19.6; Fisher’s Exact *P* < 2.2×10^−16^; **Table S5**) [6, 9].

Association of noncoding structural variants with ASD We hypothesize that rare SVs overlapping cis-regulatory elements or exons of variant-intolerant genes depleted in structural variation are associated with ASD in families. We further hypothesize that inherited SVs show a maternal origin bias, consistent with a reduced risk of ASD in females [25]. Family based association was tested using a group-wise transmission/disequilibrium test (TDT), applying it to private variants (autosomal parent allele frequency = 0.0003) assuming a dominant model of transmission [26]. To control for any potential methodological artifacts we also assessed transmission of SVs in variant-tolerant genes, which showed no transmission bias (**Table S6**). Protein coding deletions in these genes were more likely to be transmitted to individuals with ASD (54/83; transmission rate = 65.1%; *P* = 0.002), but not to controls (26/57; transmission rate = 45.6%; *P* = 0.54; Figure 2). After excluding variants that disrupted protein-coding exons, noncoding variants that intersected a predicted fetal brain promoter or UTR of a variant-intolerant gene showed a paternal transmission bias to cases (39/55; transmission rate = 70.9%; *P* = 0.0013) but not a biased maternal transmission (21/23; transmission rate = 48.9%). Controls showed a slight depletion in transmission from both parents (24/64; transmission rate = 37.5%; *P* = 0.06). The joint probability of the transmission bias in cases and controls combined was significant (joint binomial *P* = 0.003; OR = 3.2; CI = 1.2-8.7). The above associations were significant after correction for multiple testing (5 groups of SVs tested for each parent separately, **Table S6**). Validation was performed where possible using PCR, single molecule sequencing, or an in-silico SNP based approach (see methods), with a 96% validation rate overall and genotypes from validation were 100% concordant with genotype calls from SV^2^ (149/155, **Table S7**).

**Figure 2.**
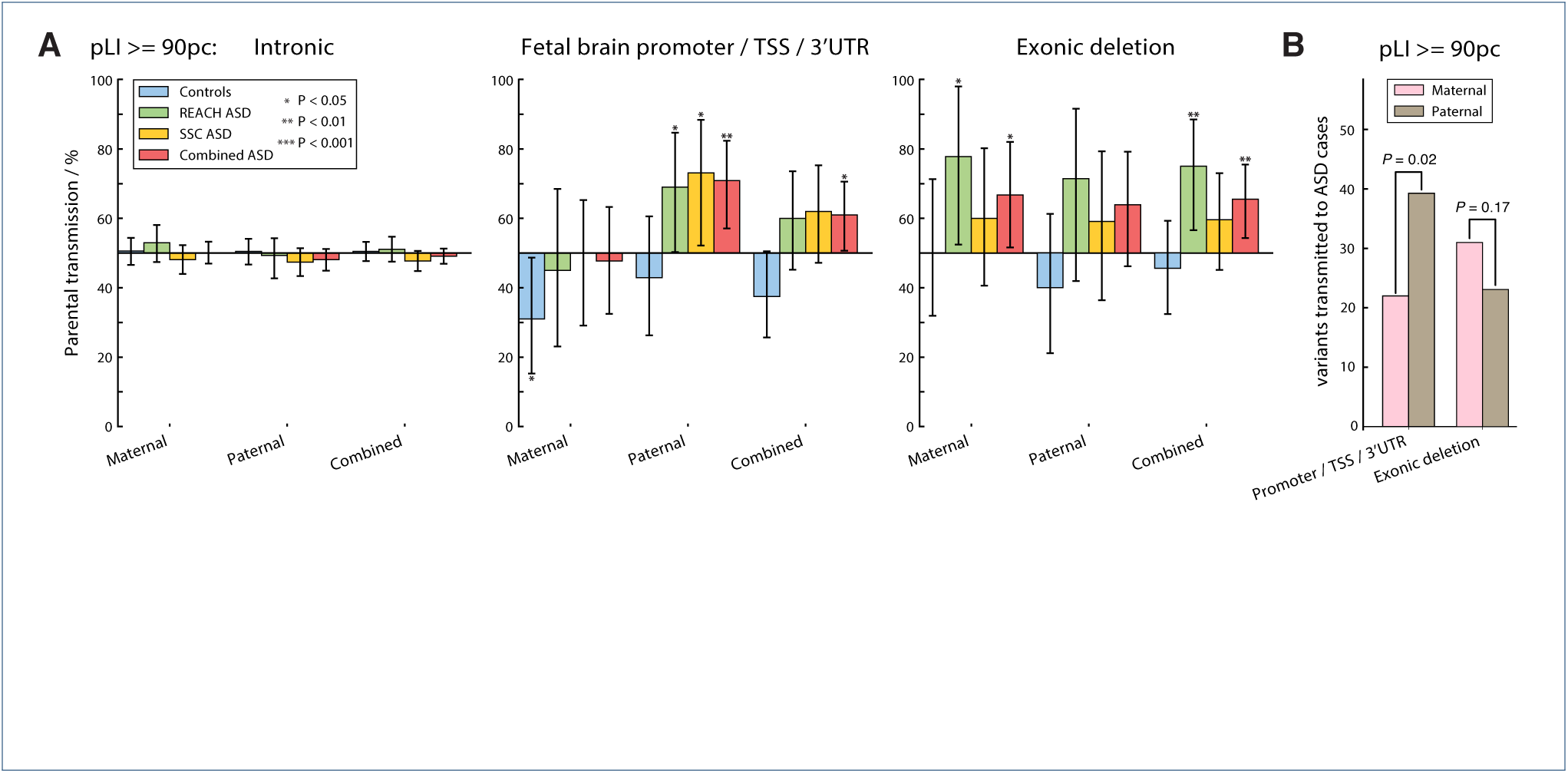
Biased transmission of noncoding and exonic variants in genes intolerant to inactivating mutations. A) Percentage of structural variants transmitted from parents to offspring that disrupt elements of variant-intolerant genes (pLI *≥*90^th^ percentile) for maternally inherited, paternally inherited, and combined parents. This analysis was stratified into controls, ASD cases from the REACH cohort, ASD cases from the SSC, and the combined ASD cases. The results reported here are for intronic variants, noncoding variants disrupting promoters / UTRs, or protein coding deletions. B) Number of variants transmitted from either the father or the mother to individuals with ASD in the combined cohort. The number of stars indicates the level of significance; whiskers represent 95% confidence intervals.

**Figure 3.**
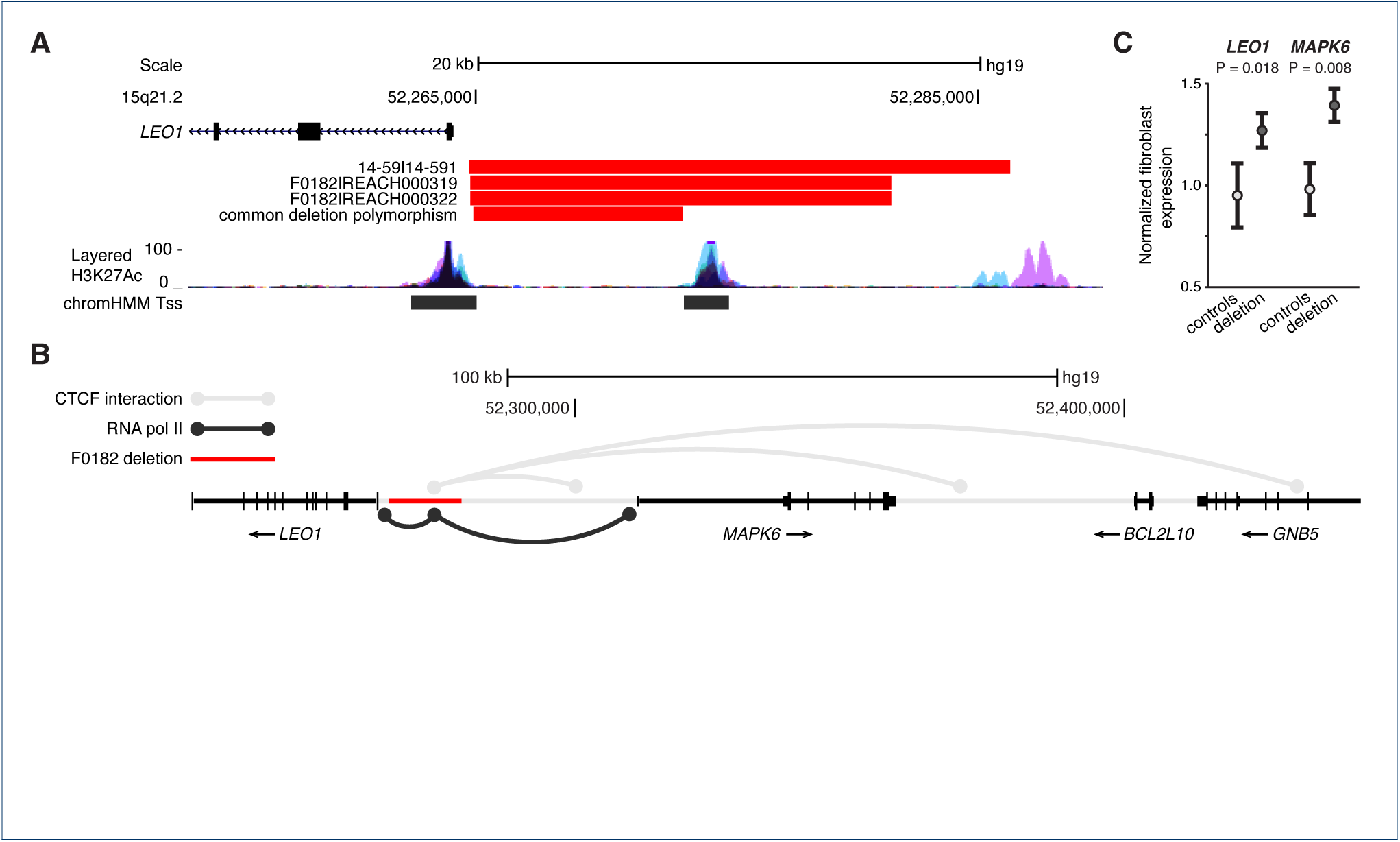
Recurrent promoter deletions of LEO1 derepress gene expression. A) Paternally inherited deletions of the *LEO1* promoter were detected in three affected individuals, one trio (14-59) and one concordant sib pair (F0182). A common deletion polymorphism (parent allele frequency = 0.011) is also present in this locus, but does not disrupt any predicted regulatory regions. B) CTCF and RNA Polymerase II ChIA-PET data suggests that the noncoding element upstream of *LEO1* disrupted by both rare deletions (F0182 deletion shown here) serves as a focal point for the spatially organized transcription of *LEO1* and *MAPK6*. C) mRNA expression of *LEO1* and *MAPK6* were increased in fibroblast lines derived from two deletion carriers (REACH000319 and REACH000322) compared to three control lines. Whiskers represent 95% Confidence Intervals. Layered H3K27Ac = Histone 3 lysine 27 acetylation (an active promoter associated mark) in seven cell types from ENCODE. ChromHMM Tss = predicted transcription start site based on chromatin signatures in multiple cell types from the Epigenomics Roadmap Project.

Further highlighting the paternal inheritance of noncoding variants, SVs in promoters or UTRs of variant-intolerant genes were more likely to be transmitted to affected offspring from the father (39 paternal, 22 maternal; Binomial *P* = 0.02; Figure 1B). A nonsignificant maternal bias was observed for coding SVs, consistent with previous studies [25, 27]. All private noncoding or protein-coding variants in genes with pLI scores ≥ 90^th^ percentile are given in **Table S7**. The median lengths of these categories of noncoding SV were 2,140bp (interquartile range IQR = 520-7,489bp) and 7,548bp (IQR = 3,795-72,664bp) respectively.

We investigated the effect of inherited SVs on autistic traits in families using the Social Responsiveness Scale (SRS) measures that were available for all family members in the SSC cohort. Parents who transmitted protein-coding deletions of variant-intolerant genes to affected probands had elevated SRS scores indicating that these variants contribute to social impairment in unaffected relatives (combined parent SV carrier mean SRS = 39.6; parent non-carrier mean = 29.1; Wilcoxon Rank Sum test *P* = 0.041; **Table S8**). Parents carrying noncoding SVs did not show significantly higher scores relative to non-carriers, and neither ASD cases nor siblings showed elevated scores in either category (**Table S8**).

Recurrent inherited gene mutations were also enriched in ASD. Five variant-intolerant genes displayed exon disrupting mutations in more than one family and were also transmitted to cases, including *ASTN2* [28], *ATAD2*, *CACNA2D3* [6], *PTPRT* and *NRXN1* [9] (**Table S7**), a 2.87-fold enrichment compared to random permutation of transmitted SVs across this gene set (expected *n* = 1.75; Permutation *P* = 0.034).

We also observed recurrent noncoding mutations in four genes, *CNTN4*, *LEO1*, *MEST* and *RAF1* (**Table S7**), a significant enrichment compared to random permutation (expected n = 0.023; Permutation P ≤ 0.0001). We examined these candidate genes in a combined dataset of 12,889 cases from 20 exome sequencing studies from ASD and developmental delay and identified two de novo mutations disrupting *LEO1* [6, 29]. This is a higher rate of *LEO1* LoF de novo mutations than would be expected based on a Poisson model that controls for gene length and sequence context (expected *n* = 0.1; *P* = 0.0025) [30, 31]. A third LoF variant of *LEO1* was reported in an ASD family [6], but parental genotypes were not available. Only one LoF mutation has been observed in this gene in 46,785 control individuals (expected *n* = 23.8) [24].

Both *LEO1* deletions eliminate an upstream regulatory element that has a chromatin signature associated with an active transcription start site in multiple cell types from the Epigenomics Roadmap Project (Figure 3) [32]. A smaller 8.7kb deletion polymorphism (parent allele frequency = 0.011) was also detected near the *LEO1* promoter, but this variant does not disrupt any annotated functional elements (Figure 3), and does not show biased transmission to cases (*P* = 0.44) or controls (*P* = 0.45). All three deletions were validated and fine-mapped by singlemolecule sequencing of long PCR products using the MinION platform (Oxford Nanopore; Figure S6).

The involvement of this functional element in gene regulation is further supported by published chromatin interaction mapped by ChIA-PET [33, 34]. Chromatin interactions associated with transcription factors CTCF and RNA polymerase II revealed this upstream cis-regulatory element to be a focal point for long range chromatin interactions associated with transcription. Expression of *LEO1* and the neighboring *MAPK6* was higher in fibroblast cell lines from two deletion carriers compared to controls (*LEO1* T test *P* = 0.018; *MAPK6 P* = 0.008; Figure 3; **Table S9**).

**Figure 4.**
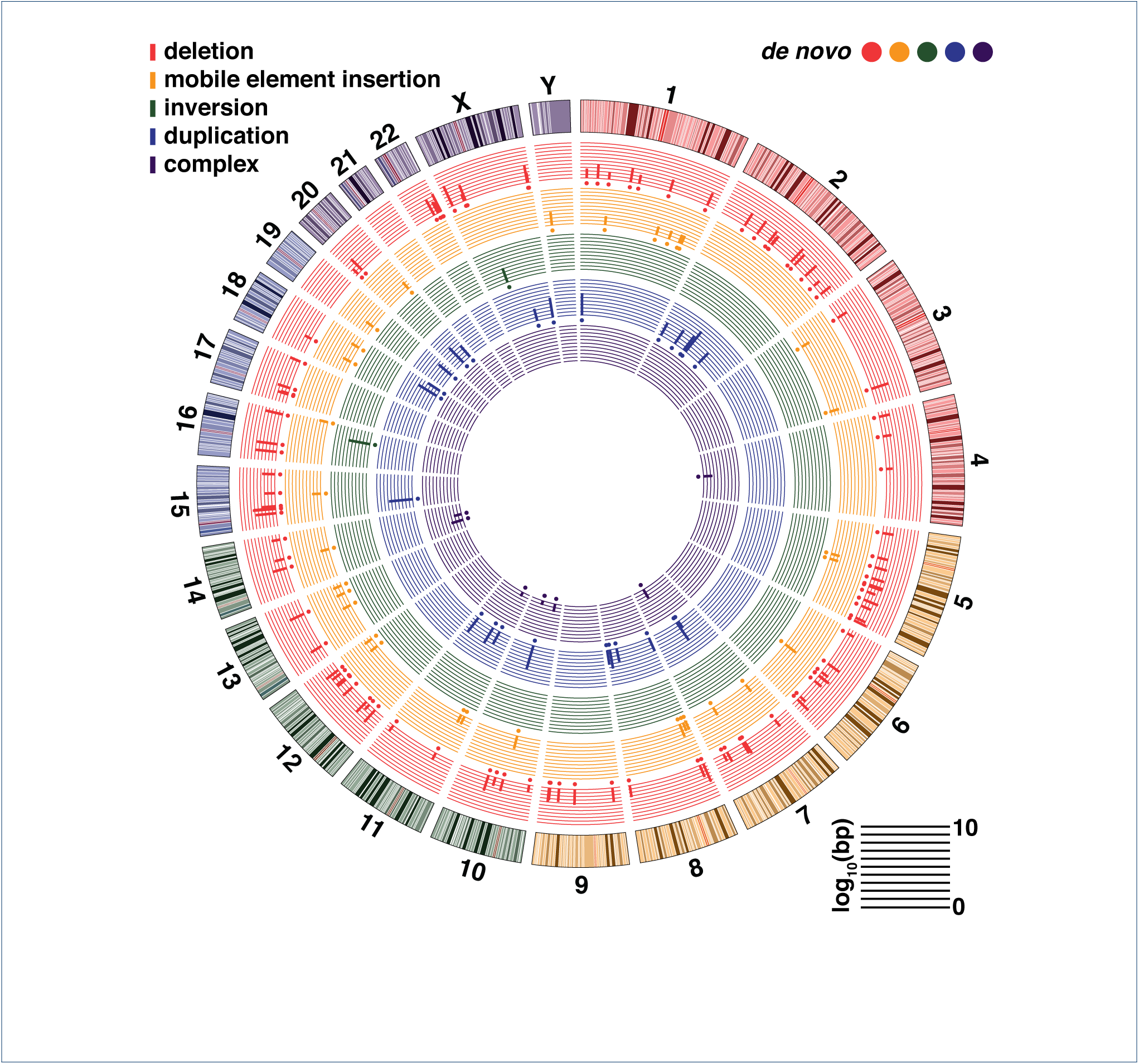
De novo structural variation in 1,510 children. Circos plot of *de novo* variants with concentric circles representing (from outermost to inner): ideogram of the human genome with colored karyotype bands (hg19), deletions, mobile element insertions, tandem duplications, balanced inversions, complex structural variants. Circles indicate the location of *de novo* SVs, and their colors match the five SV types. Bars represent the log_10_ SV length of the *de novo* variants.

**Figure 5.**
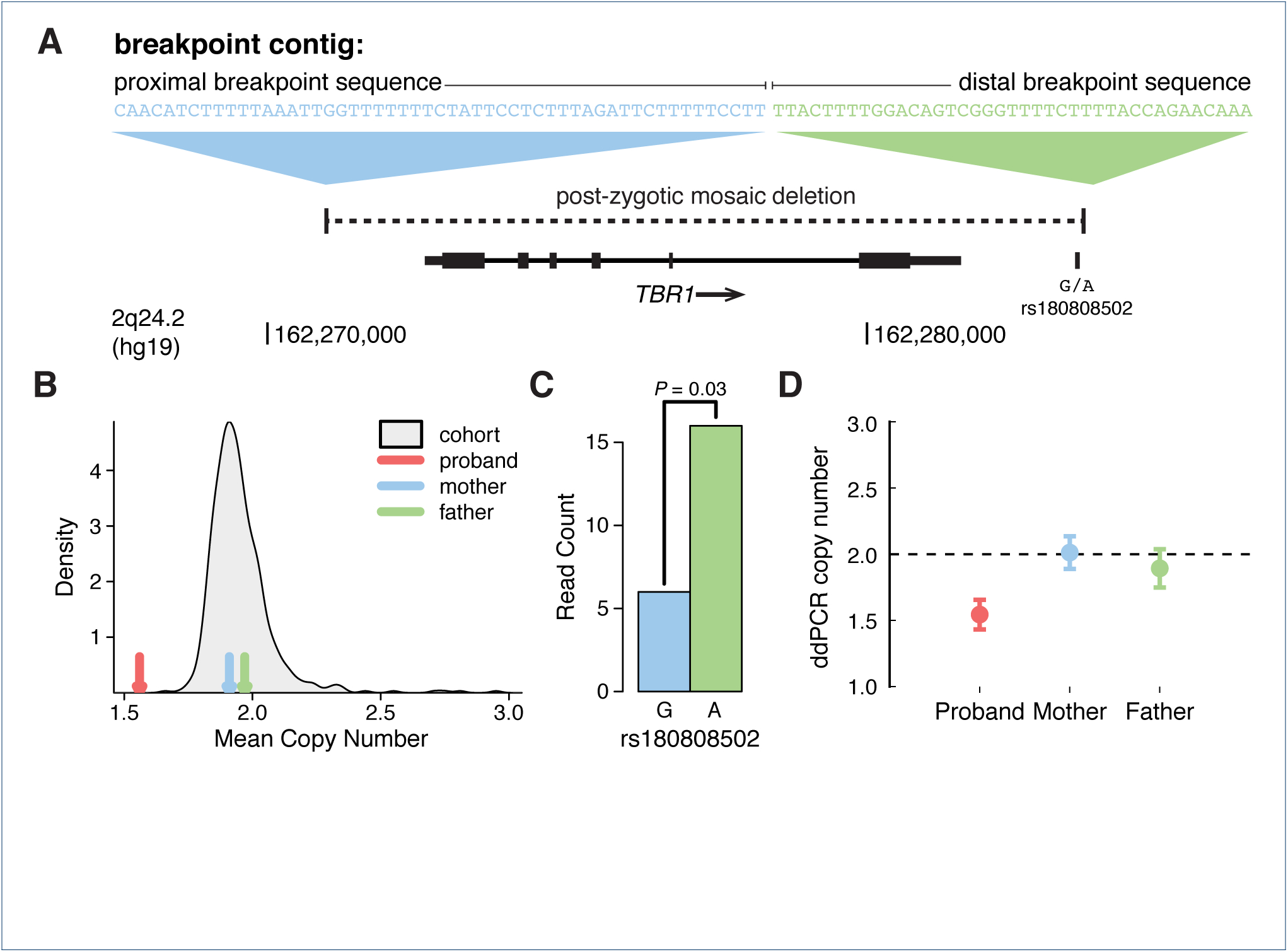
Somatic Mosaic deletion disrupting TBR1. A) Discordant paired end and split reads identify and resolve the breakpoints of a putative de novo 14.1kb deletion in the affected proband MT 18.3 that encompasses the full length of *TBR1*, but does not directly affect other genes. B) The proband’s mean coverage is intermediate between a copy number of one and two, but lower than all other individuals in the REACH cohort, indicating possible somatic mosaicism. C) There is one phased heterozygote SNP in the locus (rs180808502), which has a skewed read depth ratio indicating the deletion impacts the maternal chromosome. D) ddPCR validation confirms somatic mosaicism in the child.

### De novo and mosaic structural mutation

A circos plot in Figure 4 details the distribution of 163 de novo SVs across the genome in 1,510 children. The de novo SV mutation rate in ASD and sibling controls was 15.5% (CI = 11.8-19.9) and 12.5% (CI = 7.2-20.4) respectively in the REACH cohort (Figure S7). Despite the fact that subjects carrying de novo CNVs previously detected by microarray or exome sequencing had been excluded from the SSC sample, we detected de novo SVs in 8.7% of ASD (CI = 6.4-11.5) and 9.5% of controls (CI = 7.1-12.4; Figure S7). The FDR was 8% overall, 4.1% for variants ≥500bp (92/96 validated) and 28% for smaller variants (13/18 validated). All five false de novo variants *<*500bp proved to be false negatives in a parent (**Table S10**). All MEIs (13/13), inversions (2/2), and all but one complex SV (7/8) were validated (**Table S10**). A majority (68%) of phased de novo SVs originated from the father (binomial test *P* = 0.038; **Table S10**), similar to a previous estimate of 71% [35], and comparable to the bias observed for SNVs and indels [36, 15, 37]. Paternal age was not significantly greater in families with de novo SVs (Wilcoxon Rank Sum *P* = 0.69).

Our methods were sensitive enough to detect somatic mosaicism for a subset of deletions (*n* = 6), including a 14.1kb deletion of *TBR1* that likely occurred in the first cell division of embryonic development (Figure 5). Protein-truncating mutations of *TBR1* have been implicated as a monogenic cause of ASD [4, 38]. We estimate from our data that at least 6% (8/133; CI = 2.8-11.4%) of de novo CNVs display either high-level somatic or low-level parental mosaicism (**Table S11**), consistent with previous microarray studies [39].

Confirming what we have observed previously, de novo SNVs and indels clustered in proximity to de novo CNV breakpoints (permutation *P* = 0.0029; **Table S12**; Figure S8) [20].

### Contribution of de novo and inherited SVs to ASD

The global rate of de novo structural mutation was similar cases and controls (Figure S7), as we have previously shown [20]. The REACH cohort had a greater burden of gene disrupting de novo variants than controls (7.2% in ASD versus 2.1% in controls; permutation *P* = 9.2 × 10^−5^; **Table S13**), A 5.1% excess of gene disrupting de novo SVs in cases is slightly higher than estimates of 3-4% from previous studies using less refined methods of detection [8, 9]. A 2.3% rate of coding de novo SVs in the SSC further suggests that the contribution from small coding de novo SVs is modest (Permutation *P* = 0.46). After excluding SVs that intersected with protein coding exons, only one de novo variant-intersected a noncoding element of a variant-intolerant gene, a promoter deletion of *SIM1* in a control (**Table S10**).

Combining our findings from the REACH and SSC cohorts we are able to place lower bounds on the proportion of cases that carry rare coding and noncoding risk variants. We estimate that rare SVs contribute to 11% of ASD cases (CI = 9.8-13.4%), half of which (5.1%) are gene-disrupting de novo mutations. The remainder includes inherited rare variants of which cisregulatory and coding SVs, which contribute to 2.1% (CI = 1.2-2.8%) and 1.9% (CI = 0.8-2.9%) of cases respectively. Known pathogenic SVs not accounted for above contribute in another 1.9% of cases (**Table S14**).

## Discussion

Here we demonstrate that rare inherited SVs that disrupt cis-regulatory elements of functionally-constrained genes confer risk for ASD, and there is a similar contribution from inherited SVs that disrupt genes.

We observe a differential contribution of rare variants from mothers and fathers. SVs that disrupt variant-intolerant genes were inherited more frequently from mothers, consistent with a reduced vulnerability of females to rare variants of large effect [40, 41]. SVs that disrupt only promoters or UTRs, on the other hand, showed a significant paternal transmission bias. The underlying genetic mechanism that explains this paternal bias is not clear.

A paternal-origin effect for non-coding deletions suggests the possibility of an epigenetic mechanism. For example, deletion of key cis regulatory elements can lead to de-repression of imprinted genes [42]. Recurrent promoter deletions were observed in one gene that is known to be imprinted in fetal tissues, mesoderm specific transcript (*MEST*) [43]. However, classical [44] or brain-specific [45] imprinting are unlikely to explain our results given that both phenomenon affect a very small fraction (*<*1%) of genes and are not exclusively paternal. A paternal-specific epigenetic mechanism that acts on many functionally constrained genes has not been described, but we cannot rule out this possibility.

An alternative to an epigenetic mechanism is a ‘bilineal two-hit model’, in which risk is attributable to a combination of a maternally-inherited coding variant and a non-coding variant of moderate penetrance that is inherited from the father. Since males are more vulnerable than females to psychiatric disorders, then mothers could be more likely to carry a coding mutation of large effect [46], while additional genetic burden (including non-coding SVs) might tend to be derived from paternal lineage. A formal test of this hypothesis, however, would require a combined analysis of SNVs, indels and SVs and a much more complete knowledge of the inherited risk factors for ASD.

The intermediate genetic effects of inherited cisregulatory SVs (OR = 3.2), paternal transmission bias, and a lack of evidence for the association noncoding de novo SVs suggests that structural mutations in noncoding regions have a relatively moderate level of penetrance compared to protein coding variants. Furthermore, we demonstrate that coding variants influence neurobehavioral traits in parents, but we do not find similar evidence for noncoding variants, consistent with rare cis-regulatory variants carrying moderate risk.

SVs that directly disrupt cis-regulatory elements can identify novel candidate loci and novel genetic mechanisms underlying risk. Based on recurrent de novo LoF variants, the gene *LEO1* represents a strong candidate gene for ASD. Recurrent promoter deletions detected in this study remove a CTCF and RNA Pol II binding site that is highly topologically connected to adjacent genes, and its disruption results in the de-repression of *LEO1* and adjacent *MAPK6*.

The contribution of cis-regulatory variants that we observe was not evident in previous studies of idiopathic ASD, in part because a majority of risk SVs in this study were below the detection limits of previous methods. Furthermore, our results stand in contrast to two previous studies that have found anecdotal evidence that the rare de novo SVs of noncoding elements contribute to ASD [47, 17]. We cannot exclude the possibility that rare highly penetrant noncoding variants contribute to ASD. Indeed, there is one well-known example: the triplet repeat expansions that cause Fragile X syndrome [48]. However, we can conclude that de novo SVs within regulatory elements of variant-intolerant genes are extremely rare (observed in one control in this study).

Our analysis of distal enhancers is limited by our ability to infer the functional effects of SVs and identify their relevant target genes. Thus, it is likely that we have failed to capture some ASD risk variants in intergenic regions. A rigorous analysis of such variants would require a more comprehensive knowledge of the ‘enhancerome’ [49, 50], and an effective means for distinguishing between neutral and deleterious variants.

Due to the greater potential of SVs to impact gene function and regulation relative to SNVs and indels, this class of genetic variation has historically proven effective for illuminating new components of the genetic architecture of disease [51]. Our findings provide a demonstration of the utility of SV analysis for characterising the genetic regulatory elements that influence risk for ASD.

## Methods

### Patient Recruitment

This study consists of two primary cohorts, which will be referred to as ‘REACH’ or ‘SSC’ in the following sections. Relating genes to Adolescent and Child Health (REACH) cohort individuals were referred from clinical departments at Rady Children’s Hospital, including the Autism Discovery Institute, Psychiatry, Neurology, Speech and Occupational Therapy and the Developmental Evaluation Clinic (DEC) as part of the REACH study. Further referrals came directly through the REACH project website (http://reachproject.ucsd.edu/). In total 612 individuals from 161 families came from the REACH project. The Autism Center of Excellence at the University of California San Diego contributed 11 trios. A further 452 samples from 139 families were recruited at Hospital Universitari Mútua de Terrassa in Barcelona. The REACH families combined consisted of 112 controls and 362 affected individuals 285 with ASD, 43 with pervasive developmental disorder not otherwise specified (PDD-NOS), 10 with attention deficit hyperactivity disorder (ADHD), and 24 with speech delay, epilepsy, anxiety, or other related developmental disorders that were therefore classified as ‘cases’ for bioinformatics analyses. The Simons Simplex Collection (SSC) Whole Genome Sequencing dataset (http://bit.ly/2jc34rU) consisted of 518 quad families with sibling pairs discordant for an ASD diagnosis that were selected from the full cohort of 2,644 families [7] after excluding those where offspring carried any plausible contributory de novo or inherited SNVs, indels, deletions or duplications identified from microarray or exome sequencing data. The exclusion criteria for exome- or array-’positive’ individuals are described below and were applied to both ASD cases and sibling-controls:

1. De novo CNVs (189 families): Any confirmed or published de novo copy number variant (CNV) [52, 53], Illumina SNP genotyping data, or exome CNV data that is: Rare (≤0.1 population frequency based on parents and DGV) or genic (≥1 exon).
2. Inherited CNVs (92 families): Any CNV from Illumina genotyping data [53], or exome CNV data that is: rare (≤0.1 population frequency based on parents and DGV), or intersects ≥10 genes.
3. De novo LoF (564 families): Any de novo loss of function from published sequencing data that is: rare (≤0.1 population frequency based on the exome variant server), nonsense, canonical splice site, or frameshift [7, 40].

### Whole Genome Sequencing

Our combined dataset consisted of WGS data collected for two cohorts and sequenced at three sites (**Table S1**). All WGS data were generated from whole blood DNA. All members of individual families were sequenced within the same batch of samples.

### REACH cohort

The REACH cohort initially consisted of 1,126 individuals from 319 families, including 893 individuals from 260 families that were sequenced at Human Longevity Inc. (HLI) on an Illumina HiSeq X10 system (150 bp paired ends at mean coverage of 50X) and an additional 204 individuals from 59 families that were sequenced at the Illumina FastTrack service laboratory on the Illumina HiSeq 2500 platform as described in our previous publication [20]. We performed initial quality control (QC) steps to ensure relatedness and gender matched the sample sheets, excluding any mismatches or half-siblings. We also tested for an excess of Mendelian errors in the children, and an excess of single nucleotide variants called in either parent (≥3 SD from the mean) indicative of low quality DNA. In total 29 samples were removed, including eight complete families. Therefore, 1,097 individuals from 311 families were taken forward for structural variant calling and analysis.

### SSC Cohort

Whole genome sequencing of the SSC cohort on an initial 540 families was performed at the New York Genome Center on an Illumina HiSeq X10 (150 bp paired ends at mean coverage of 40X). Of the 540 SSC families, 518 were complete quad families. Incomplete families were excluded from the dataset. All 518 met the above QC criteria for inclusion in the study. Mean coverage (39-50X) and insert sizes (348-420) and were similar at all three sequencing sites (**Table S1**). Sequence alignment and variant calls for REACH samples were generated on families using our WGS analysis pipeline implemented on the Comet compute cluster at REACH. For SSC samples the same pipeline was adapted for use on Amazon Web Services (AWS). In brief, short reads were mapped to the hg19 reference genome by BWA-mem version 0.7.12 [54]. Subsequent processing was carried out using SAMtools version 1.2 [54], GATK version 3.3 [55], and Picard tools version 1.129, which consisted of the following steps: sorting and merging of the BAM files, indel realignment, removal of duplicate reads, base quality score recalibration for each individual [56].

### SV Detection

We utilized four complementary algorithms to detect SVs: ForestSV, Lumpy, Manta, and Mobster. ForestSV is designed to detect deletions and duplications based on a combination of signatures including, coverage, discordant paired ends and other metrics such as mapping quality [15]. In addition we implemented two algorithms, Lumpy and Manta (Manta workflow version 0.29.0 was run with default parameters), the latter being a new addition to the SV analysis pipeline since our previous publication [20], both of which utilize a combination of discordant paired ends and split reads and have greater sensitivity for small (*<*500 bp) deletions, duplications, inversions and complex rearrangements [57, 58, 59]. Finally, Mobster uses discordant paired-end and split-read signal in combination with consensus sequences of known active transposable elements to identify mobile element insertions (MEIs) [60]. A consensus callset was generated by merging calls from ForestSV, Lumpy, Manta and Mobster. SV calls from multiple methods were combined, and overlapping variants detected in the same sample were collapsed as described in our previous structural variant publication [20]. The unfiltered consensus callset consisted of the union of calls from the four methods. As a preliminary filtering step, SVs were removed from the consensus callset if they overlapped by more than 66% with centromeres, segmental duplications, regions with low mappability with 100bp reads, regions subject to somatic V(D)J recombination (parts of anitbodies and T-cell receptor genes). SVs called by Manta or Lumpy were filtered if they had one or both breakpoints overlapping one of these regions. Regions used for filtering can be found in our previous publication [20].

### SV genotyping and filtering

We generated a set of uniformly-called genotypes for the combined set of deletions and duplications called by three methods Lumpy, Manta, or ForestSV, using a single genotyping algorithm SV^2^ v2.0 (https://github.com/dantaki/SV2). SV^2^ provides estimates of genotype likelihoods for deletions and duplications across a broad size range (10bp-10Mb), and this metric was used as our primary filtering criterion for these. The SV callers Lumpy [58] and Manta [59] provide genotype likelihoods for the subset of calls that were generated by these methods, which include SVs that are not genotyped by SV^2^ such as inversions and nontandem duplications. These genotype likelihoods were also used as quality metrics during the filtering of SV callsett as described below.

We assessed the performance of each genotyper for deletions and duplications across a range of sizes and depending on sequence context (short tandem repeats, segmental duplications, etc.), estimating the FDR from Illumina 2.5M SNP array data on a subset of 205 genomes using the Intensity Rank Sum test implemented using the Structural Variation Toolkit. Based on these FDR estimates, we applied a range of genotype likelihood filters on variants. For de novo SV calling, more stringent SV^2^ genotype likelihood filters were applied to safeguard against false positives in the child or false negatives in the parents, including a minimum reference genotype likelihood. The final filtering criteria are detailed in **Table S3**.

Genotype-likelihood thresholds for SV filtering were determined based on estimates of FDR, which were performed from Illumina 2.5M SNP array data on a subset of 205 genomes using the Intensity Rank Sum test implemented using the Structural Variation Toolkit. SV^2^ designates SV calls as ‘PASS’ or ‘FAIL’ at two levels of stringency: ‘standard’ and ‘de novo’, which are described in detail in our companion paper [23]. Standard filters were used to generate to overall callset and for family based association testing. The more stringent de novo filters were used for de novo mutation calling. In addition, we included in the consensus callset SVs, which passed genotype likelihood thresholds for Lumpy and Manta, and thresholds were selected based on FDR estimates for SVs across a range of sizes and depending on sequence context (short tandem repeats, segmental duplications, etc.). FDR estimates for SV calls filtered at standard and de novo stringency and genotype likelihood thresholds for Lumpy and Manta are provided in **Table S3**.

Due to the requirements of this study for high genotyping accuracy, we have applied additional filtering measures that were not used in a previous publication from our group [20]. The FDR of variants intersecting STRs was an order of magnitude higher than SVs that did not; therefore more stringent genotype likelihood filters were applied to SVs overlapping STRs (**Table S3**). Furthermore because STRs were depleted in probes on the Illumina 2.5M SNP array, only 7.2% of deletions and 12.9% of duplications overlapping an STR had one or more probes, compared to 28.5% of deletions and 56.3% of duplications that do not. FDR estimates for these variants could be less accurate. Therefore, for all analyses in this study, we have excluded SVs with breakpoints overlapping STRs. We have also annotated these in the callset VCF (which can be downloaded from NDAR study number 434), and we suggest that these SVs be treated with caution. Hence, the number of deletions and duplications reported in the SV callset here is lower than in our previous publication [20].

Deletions and duplications called by Lumpy and Manta were overrepresented by breakpoints that overlap with short tandem repeats (STRs) 21.75 and 49.6% respectively compared to the 2.3% of the genome that consists of STR. The FDR of variants intersecting STRs was also an order of magnitude higher than SVs that did not; therefore more stringent genotype likelihood filters were applied to SVs overlapping STRs (**Table S3**). Furthermore because STRs were depleted in probes on the Illumina 2.5M SNP array, only 7.2% of deletions and 12.9% of duplications overlapping an STR had one or more probes, compared to 28.5% of deletions and 56.3% of duplications that do not. FDR estimates for these variants could be less accurate. It is therefore suggested that these SVs be treated with caution (they are annotated in the callset VCF, which can be downloaded from NDAR study no. 434). We have excluded SVs with breakpoints overlapping STRs for all analyses. Due the high stringency filters that were applied to this subset of variants, the number of deletions and duplications reported in the SV callset here is lower than in our previous publication [19, 20].

In total we detected 11.87 million alleles from 89,123 distinct loci encompassing 19.4% of the GRCh37 (hg19) release of the ‘mappable’ reference human genome (0.497/2.57Gb, excluding SVs larger than 1Mb, which are likely to be pathogenic and would contribute disproportionately to this estimate, **Table S2**). 12.5% (320Mb) of the reference genome was deleted and 7.3% (186Mb) duplicated in our cohort of 829 families.

### De novo calling and phasing

De novo SVs were called if they occurred in a child and were genotyped reference in both parents and the parent allele frequency for the variant was less than 1%. We also applied more stringent SV^2^ genotype likelihood filters for de novo SVs and TDT analyses, which are detailed in **Table S3**. The average rate of Mendelian errors in the callset as a whole for deletions and duplications was 0.99% (95% CI: 0.03) and 4.66% (95% CI: 0.15) respectively (Figure S4). De novo genotype likelihood filters applied to variants with parent allele frequencies *<*1% reduced the rate to 0.21% (95% CI: 0.1) for deletions and 0.5% (95% CI: 0.2) for duplications.

### SV validation

We validated large putative de novo deletions and duplications using an in silico SNP-based approach that utilizes read depth from the VCF files from GATK Haplotype Caller. For each SNP we normalized allelic read depth relative to the genome average for reference / alternate alleles, and calculated a z-score for each SNP. We also calculated the B allele frequency (BAF) by taking the average of the allele (reference or alternate) with the fewest number of supporting reads across the locus. Since deletions are hemizygous the expected BAF is 0 (unless the mutation is mosaic, see below). For duplications we calculated the BAF only for heterozygote SNPs, which have an expected BAF of 0.33 for autosomal variants. If the child showed an average elevated or depleted SNP read depth more than one standard deviation from both parents, and a BAF consistent with the called SV type, and / or the variant could be phased, then the SV was designated as valid. Furthermore this SNP data was used to determine the parent of origin, by performing a paired t-test on phased SNP allelic depth within the locus. We plotted the validation results for each member of the trio using the R package CNVplot, which was developed in house (https://github.com/dantaki/CNVplot). The plots can be viewed by clicking on hyperlinks in **Tables S7, S10, and S13**. This approach is orthogonal to the SV calling steps above, which do not phase variants, calculate their BAF, or estimate coverage using SNP data.

Small deletions, duplications, inversions, complex SVs, and MEIs were validated using PCR. Both de novo inversion calls were validated. We attempted PCR validation on 13 de novo *Alu* elements, all of which validated as de novo. *Alu* insertions have polyA tails; we therefore used a lower extension temperature (65°C), because A/T rich sequences have a low melting temperature [61]. We also used longer extension times (90 seconds) to an otherwise standard PCR protocol.

### Oxford Nanopore Validation

Recurrent deletion of the *LEO1* locus were validated and fine mapped by single molecule sequencing. Deletions and wild type sequence were amplified by long range PCR in three families with *LEO1* deletions (14-59, F0182, and F0208). We performed reactions in a volume of 10*μ*l PCR, containing contained 20ng of patient genomic DNA, 0.4*μ*M forward and reverse primers and LongAmp^®^ Taq 2X Master Mix (New England BioLabs, M0287L). We gel-purified PCR amplicons and barcoded them using Oxford Nanopore Technologies’ (ONT) Native Barcoding Kit 1D (EXPNBD103) and added sequencing adapters using Ligation Sequencing Kit 1D (SQK-LSK108). We ran sequencing libraries for 48 hours on ONT’s MinION Mk1B, using the SpotON Flow Cell Mk I (R9.4, FLOSPOTR9) and MinKNOW software (v.1.3.30). In total, we generated approximately 2.3Gb of fasta data. We applied a quality and length filter was applied to the unaligned reads and removed those with a mean quality score of 8.5 or less, or which differed from the expected amplicon length by 2kb or more. Using BWA-mem (v.0.7.15-r1140) [54] with the ‘-x ont2d -M’ flags we aligned reads to the human genome (hg19), and filtered to keep those that overlapped the amplicon region. Regions of high coverage were defined as those areas where the coverage was 20% or higher of the maximum coverage for that amplicon. For each of the deletion amplicons, we analysed the coverage profile to determine putative deletion endpoints, and used these endpoints to generate a putative haplotype sequence using the reference genome. We also generated a corresponding wild-type haplotype. We realigned reads using BWA-mem against these haplotypes and then filtered read that did not align to the expected haplotype or that covered less than 95% of the high coverage regions. We fed the alignments for the top 100 reads, as judged by read quality score, into nanopolish (v.0.6-dev, commit 8be00b94182, https://github.com/jts/nanopolish/) [62] to generate a consensus, and called SNPs using Mummer [63]. The consensus fasta sequences can be downloaded from NDAR.

### Evaluation of SV calling across data from multiple sequencing centers

The average SV numbers for each class of SV were similar between cohorts sequenced at different sequencing centers (**Table S1**). We compared SV calls for one individual (REACH000236) who was sequenced twice, on the Illumina HiSeq 2500 with 100bp reads (at 43X coverage) and on the Illumina HiSeq X with 150bp reads (also at 43X coverage). Since the coverage is the same between the two samples but the read length is 50% longer on the HiSeq X, this sample has only 2/3 as many reads when sequenced on the HiSeq X. This affects SV calling for two reasons, there will be on average more split reads supporting each call on the HiSeq X, but fewer discordant paired-end reads. The overlap between the SVs called on each platform in this sample ranged from 66-96% for each SV type (Figure S9).

### Investigating the intolerance of genetic functional elements to structural variation

We investigated the enrichment/depletion of private deletions, duplications, and mobile element insertions within specific genomic features compared to a random distribution of SVs, we shuffled the position of sites that were private to families (i.e. observed in only one parent) across the genome using BedTools [64], while excluding overlap with regions of the genome that cannot be sequenced with short reads. We counted the number of times where a shuffled SV overlaps (at least 1bp) the following genomic features: protein coding exons, transcription start sites (TSS), 5’UTRs, 3’UTRs, promoters, noncoding RNAs, enhancers, conserved noncoding regions, human accelerated regions, CTCF binding sites, exon flanking (one breakpoint within 100bp of an exon), 1kb upstream, 1kb downstream, and introns. Events that overlapped multiple features were prioritized in the order above, so for example if a variant overlapped a protein coding exon, a 3’UTR and an intron, it is counted as protein coding but not 3’UTR or intronic. Each feature is explained in detail below and we’ve summarized each in a table included as part of **Table S4**. We performed 10,000 permutations and compared the observed overlap to the expected overlap. *P* values were corrected using a Benjamini-Hochberg false-discovery rate adjustment.

### Definitions of gene disrupting SVs versus noncoding

Gene disrupting deletions were defined as those that directly disrupted at least one protein coding exon from one transcript of a gene (transcripts were extracted from hg19 refgene). Noncoding deletions could delete UTRs, introns, enhancers, or promoters of genes, but not protein coding exonic sequence or the start position of the first exon of a transcript. Protein coding duplications were divided into four categories. Whole gene duplications encompassed at least one full length transcript of a gene. Internal exon duplications intersected at least one protein coding exon internal to a transcript, but not the UTRs. Duplications that intersected at least one exon and with one breakpoint outside of the gene and the other internal to the gene were divided into two categories, those that encompassed the 5’UTR (and promoter), and those that encompassed the 3’UTR. Gene disrupting inversions were classified as variants that either had one or both breakpoints inside a protein coding exon of a gene, or that had one breakpoint in an intron of a gene and the other breakpoint either outside of that gene or in another intron. Inversions that inverted an entire gene or genes but had intergenic breakpoints were considered noncoding.

### Definition and selection of noncoding elements

Transcription start sites, 3’UTRs, and 5’UTRs were defined using full-length protein-coding transcripts from RefSeq. We defined two types of noncoding RNAs, micro-RNAs and natural antisense transcripts. Human micro-RNAs were downloaded from miRBase (v21) [65], lifted over to hg19 annotated to genes if they were intronic in a sense orientation and therefore transcribed with the gene itself. We assigned exons of natural antisense transcripts (NATs) to genes if they were transcribed in an antisense direction and overlapped with a gene. NAT data was downloaded from GENCODE v25 (only including transcripts with support level of 1, 2 or 3) [66].

Conserved noncoding regions were defined from two studies; one that defined ultraconserved elements ≥100bp conserved in human, mouse and rat genomes [67], and the other that defined ultrasensitive noncoding regions with almost as much selective constraint as coding genes [68].

We defined promoters and enhancers using fetal brain data Epigenomics Roadmap Project and data from ENCODE [32]. The Epigenomics Roadmap Project integrated combinatorial interactions between five different chromatin marks to define 15 chromatin states using a Hidden Markov Model algorithm called chromHMM v.1.10 [69] (http://egg2.wustl.edu/roadmap/web_portal/chr_state_learning.html). Four states were used to define promoters, active transcription start site (1 TssA), TSS flank (2 TssAFlnk), bivalent TSS (10 TssBiv), and bivalent TSS flank (11 BivFlnk). Three states were used to define fetal brain enhancers, genic enhancer (6 EnhG), enhancer (7 Enh), and bivalent enhancer (12 EnhBiv). For the Epigenomics Roadmap Project data we defined fetal brain promoters/enhancers using the intersection of male and female fetal brain tissue (epigenomes: E081 and E082). We defined adult brain promoters/enhancers using the intersection of epigenomes from eight brain regions (E067 (Angular gyrus), E068 (Anterior Caudate), E069 (Cingulate Gyrus), E070 (Germinal Matrix), E071 (Hippocampus), E071 (Inferior Temporal Lobe), E073 (Dorsolateral Prefrontal Cortex), and E074 (Substantia Nigra)), excluding any elements that intersected with those in fetal brain.

ENCODE enhancers and promoters were defined based on chromatin state segmentations from six human cell lines (GM12878, K562, H1-hESC, HeLa-S3, HepG2, and HUVEC), which integrated ENCODE ChIP-seq, DNase-seq, and FAIRE-seq data from two algorithms (chromHMM and Segway) to segment the genome into seven states [69, 70]. Data for all six cell types was downloaded from UCSC genome browser, two states were used to defined ENCODE promoters, predicted promoter or transcription start site (state: TSS), predicted promoter flanking region (state: PF). One state was used to define ENCODE enhancers, predicted strong enhancer (State: E). ENCODE CTCF enriched elements were used to define CTCF binding sites (State: CTCF). Promoters and Enhancers were assigned to genes based on proximity, if they intersected or were within 10kb of the transcription start site of an isoform of the gene.

Assigning enhancers to genes based purely on proximity is not the most effective approach, as the majority of annotated enhancers do not interact with the nearest gene [71, 50]. We therefore implemented TargetFinder, a machine-learning algorithm that annotates to genes with an FDR ≤15% by integrating features such as DNA methylation, histone marks, and cap analysis of gene expression (CAGE) data to predict distal enhancers (distance 10kb-2Mb) that interact with promoters [50]. We extracted all enhancers predicted to directly activate genes in six cell types from ENCODE (GM12878, HeLa-S3, HUVEC,

IMR90, K562, and NHEK) [50]. We also attempted to assign enhancers to genes using the correlation of expression between enhancers and promoters within 500kb of each other using data from FANTOM5 [49].

We downloaded chromatin interaction analysis by paired-end tag (ChIA-PET) data detailing the interactome map between noncoding elements and transcription start sites through CTCF or RNA polymerase II interactions [33, 34]. For each interacting pair of elements if one member of the pair overlapped a promoter of a gene (within 10kb) we assigned its pair to the target gene as a putative noncoding interacting element. Finally we also tested fetal central nervous system DNase hypersensitivity data [17] and ‘human accelerated regions’ that have undergone rapid evolution since the split from chimpanzees [47]. Both these features were assigned to genes based on proximity as for enhancers and promoters.

### Defining variant-intolerant genes and annotating known ASD genes

We categorized genes based on their probability of being loss-of-function (LoF) intolerant (pLI) as assessed by large-scale exome sequencing of populations by the Exome Aggregation consortium (ExAC) [24]. We downloaded the data from EXAC release 0.3.1 (January 2016), and used the scores calculated using a subset of the data that excluded individuals with schizophrenia. The pLI score ranges from 0-1 for 18,421 genes, with higher scores indicating that a gene is more intolerant to inactivating mutations.

Our set of known autism genes were taken from the integration of ASD array data and exome sequencing of the SSC cohort [9], and genes with an FDR ≤0.1 from another large scale whole exome sequencing study [6]. In total there are 71 ASD associated genes.

### Transmission Disequilibrium Test

For family-based association tests, we used SV^2^ genotype calls for SVs filtered at standard stringency. We tested whether variants private to families in our callset were transmitted to affected children or controls more or less than expected by chance, using a twotailed haplotype-based group-wise transmission disequilibrium test (gTDT) [26], assuming a dominant model. We excluded variants smaller than 100bp or overlapping STRs (≥50%) as it is challenging to validate them or estimate their FDR. We further excluded two families from this analysis, one family where the parents DNA was cell line derived (MT 121), and one family where the mother and child had an excess of coverage based calls from ForestSV (F0226). Our analysis focused on genes with pLI scores ≥90^th^ percentile, which we determined are enriched for genes associated with autism from published exome studies. We also only tested features that were depleted in structural variation from the callset permutation analyses above as we hypothesize that these features will be enriched for variants associated with autism.

*P* values were corrected for multiple testing using a Benjamini-Hochberg false-discovery rate adjustment, and both the coding and noncoding results detailed in the main text pass a false discovery threshold of 1%.

To compare paternal and maternal transmission rates to cases we performed a binomial test under the assumption that 50% of transmitted variants should derive from each parent. Case-control transmission analyses were performed using a joint-probability binomial test, by combining transmission of both cases and controls into a single association test. We defined association supporting transmission events as those that were transmitted to cases or untransmitted to controls, and transmission events not supporting association as those that were untransmitted to cases or transmitted to controls. We then performed a binomial test on these two groups to calculate the joint probability.

### Considering potential biases or technical artifacts in the TDT

The transmission disequilibrium test requires accurate genotyping of variants. Genotyping error can result in the apparent biased transmission of parental variants to offspring. For example false-positive SV calls in parents or false negative genotype calls in children can lead to an apparent under-transmission bias. For instance, given an FDR of 2% for SV calls in parents, and no transmission of the false calls, a rate of 48% transmission would be consistent with random segregation. This modest under-transmission bias, is not specific to SVs, and is also apparent for single nucleotide variants genotyped using GATK [26]. Ascertainment bias for rare SVs could potentially have similar effects. For example, families with many children could be prone to an overtransmission bias because variants present in parents and multiple offspring could be better ascertained than untransmitted variants present in only one parent.

We have therefore evaluated the potential for genotyping error to lead to spurious results in the TDT as part of a companion study [23] and in this study, we further examined the rates of Mendelian error and transmission to offspring for private SVs across a broad size range (Figure S4). Our results suggest that private *>*100 bp deletions and duplications respectively have low FDR (2.3% and 1.7%) and Mendelian error rates (2.0% and 0.6%). As expected based on the 4% FDR for deletions 100bp-1kb, there is a subtle (2.0%) undertransmission bias, which is consistent with random segregation of these variants (Figure S4). Since only 2.7% of variants *<*100bp had probes on the Illumina 2.5M SNP microarray we could not accurately estimate the FDR; therefore these SVs were not included in our analysis.

Our development of a machine-leaning genotyping algorithm, SV^2^, has enabled us to obtain genotype calls with high accuracy, thus eliminating such bias for SVs [23]. As expected based on the FDR, there is a subtle (1-2%) undertransmission bias for variants *<*1kb (**Table S6**), No bias is apparent for SVs ≥ 1kb (**Table S6**).

As an additional control in the TDT we also demonstrate that there is no transmission bias for intronic variants (which are not depleted in SVs), and we tested all features in genes with pLI scores <90^th^ percentile. Both ‘control’ sets of SVs were suitable as comparators as they did not differ in terms of SV length, family-size or genotype likelihoods of SVs in functionally constrained genes. We were therefore able to rule out a systematic transmission bias as an explanation for our results. Lastly, over-transmission of private coding and non-coding SVs was specific to cases, not observed in controls, and the association was replicated in an independent cohort.

### Permutations of recurrent SVs

To permute the relative enrichment / depletion of SVs overlapping the same functional elements (e.g. exons) in different families, we permuted these variants across the genome ensuring that permuted variants intersected at least one functional element of a gene with a pLI score ≥90^th^ percentile using bedtools shuffle (by implementing the -incl command). For analysis of coding variants we required that observed / permuted variants hit any exon of the same gene to be considered recurrent. For noncoding analysis we required that variants hit the same element (e.g. a 5’UTR from the same transcript) to be considered recurrent. We counted the number of times we observed a gene or functional element was intersected by more than one distinct SV and compared this to 10,000 permutations.

### Testing the association of *LEO1* de novo mutations with ASD and DD

A series of 20 different studies have been published that reported all de novo mutations detected across the exome in cases. For a specific candidate locus in this study we have investigated the potential association with developmental disorders base on tests of de novo mutation burden in a large combined sample of 13,391 subjects.

### SV Burden

The burden of de novo structural variants between individuals with ASD in this study and the controls from this study was assessed using a case-control permutation test implemented in PLINK [72].

### Parental Mosaic Structural Variation

If one parent was genotyped as ‘reference’ by SV^2^ but had intermediate copy number estimates and / or low levels of discordant paired-end / split read support for the de novo variant, we considered them to be potentially mosaic in that parent. We therefore validated all of these variants with PCR and Sanger sequencing and then estimated the levels of parental mosaicism using a custom designed ddPCR assay with a FAM labeled probe that spanned the breakpoints, and a HEX labeled RPP30 reference assay (BioRad laboratories). We assessed the copy number of the deletion breakpoint in the child, the putative mosaic parent, and the other parent as a negative control.

### Post-Zygotic Mosaic Structural Variation

We estimated the copy number of de novo copy number variants using SV^2^, and if a de novo deletion showed intermediate copy numbers (i.e. between 1 and 2) and the BAF was consistent with heterozygosity within the deletion region, this is suggestive of somatic mosaicism. We therefore phased heterozygous SNPs and determined if paternal or maternal alleles had consistently lower or higher allelic depth by performing a paired T-test (or a binomial test in the case where there was only one phased SNP). Standard copy number estimating ddPCR assays (BioRad Laboratories) were performed to validate mosaics.

### Mutational Clustering

To assess whether de novo SVs cluster with de novo nucleotide substitutions or indels, we used a window based permutation approach. We took windows of 100bp, 1kb, 10kb, 100kb, 1Mb, and 10Mb around the breakpoints of de novo SVs and intersected the windows with de novo SNVs and indels in the same individuals (de novo detection of SNVs and indels was performed as described in our previous publication [20]). We then shuffled the position of these windows in the genome either randomly (excluding regions that were filtered during SV calling) or across detected inherited SV breakpoints using BedTools and calculated the expected number of window overlapping DNMs using 100,000 permutations.

### Overlap of Structural Variants with known regions associated with developmental disorders

CNV regions associated with autism or schizophrenia were taken from three large studies, detailed in **Table S7** [8, 9, 73].

### Fibroblast cell culture and quantitative RT-PCR

Dermal fibroblasts were obtained from the California Institute for Regenerative Medicine (CIRM) (Oakland, CA, USA) or obtained from N. Chi (University of California, San Diego). Samples used for analysis included fibroblasts from F0182 | REACH000322 (ASD proband and deletion heterozygote), F0182 | REACH000321 (father, deletion heterozygote), and three unrelated control samples: CW60038, CW60044, and JS034. Cells were recovered from cryogenic storage as per CIRM’s protocol and cultured in Dulbecco’s modified eagle medium (DMEM) supplemented with 10% fetal bovine serum, 2 mM L-glutamine, 100*μ*g/ml penicillin and 100*μ*g/ml streptomycin (Thermo Fisher Scientific, Waltham, MA, USA). Cells were maintained in an incubator at 37°C at 5% CO2 and harvested for RNA isolation at passage three.

Total RNA was isolated using the Quick-RNA Microprep kit (Zymo Research, Irvine, CA, USA) protocol for adherent cells with in-column DNAse treatment. cDNA was synthesized from 100ng of RNA using random oligo primers as part of the High Capacity cDNA Reverse Transcription kit (Applied Biosystems, Foster City, CA, USA) according to the manufacturer’s protocol. Multiplexed qPCR reactions were conducted in triplicate for each sample using gene-specific predesigned PrimeTime^®^ qPCR assays for *LEO1* (Hs.PT.58.448164, FAM-labeled) and the housekeeping gene *HPRT1* (Hs.PT.58v.45621572, HEX-labeled) (Integrated DNA Technologies, Coralville, IA, USA) on a CFX Connect Real-Time PCR System (Bio-Rad, Hercules, CA, USA) along with no-template and no-reverse-transcription controls. Changes in gene expression were calculated using the comparative CT method [74] and the null hypothesis was assessed using a Student’s two-tailed unpaired T-test.

## Author’s contributions

Conceptualization, J.S., W.M.B; Methodology, W.M.B., D.A., M.G., J.S.; Software, D.A., W.M.B., M.G.; Validation, M.M., T.R.C., S.T., M.L.K., Y.Y., E.H.; Formal Analysis, W.M.B, D.A., M.G., M.L.K., P.T., K.S.M.; Writing – Original Draft, W.M.B., J.S.; Writing – Review and Editing, L.M.I, A.M., D.J.T., C.M.N.; Resources, K.K.V., T.P., S.C.T., B.K., A.T., J.C.V, C.C., N.A., A.R.M., R.C., B.C., L.M.I., A.H., M.J.A., I.R., S.J., D.J.T., S.F.K., J.G.G.,E.C.,K.P.; Visualization, W.M.B., D.A.; Supervision, J.S.; Project Administration, O.H.; Funding Acquisition, J.S., W.M.B., D.A., M.K.

## Acknowledgements

We would like to thank the families who volunteered for the study. We would also like to thank Wayne Pfeiffer, Mahidhar Tatineni, Amit Majumdar, Shawn Strande, the San Diego Supercomputer Center, and Amazon Web Services for hosting the computing infrastructure necessary for completing this project. This study was supported by grants to JS from the NIH (MH076431) and Simons Foundation Autism Research Initiative. Further support to J.S. and K.K.V. from the ASD enlight foundation. Postdoctoral Fellowships to WB from the Autism Science Foundation and MLK from the Canadian Institutes of Health Research. A T32 training grant to DA from the NIH. A.R.M. is supported by a NARSAD and NIH grants R01MH108528 and R01MH109885. Funding for K.P. is from the NIMH (R01MH110558). Funding for B.C. is from MINECO (SAF2015-68341-R), AGAUR (2014-SGR-0932), La Marató de TV3 (092020), and the European Comission H2020 Programme MiND (643051). Funding for E.C. is from the NIMH (R01MH110558, I-P50-MH081755) and a Simons Foundation Grant. L.M.I. was supported by NIH (R21 MH104766, R01 MH105524, and R01 MH109885) and in part by the Simons Foundation grant (345469). A.H. and M.J.A. received grant support from the Institute Carlos III (FIS PI11/00620) and Mutua Terrassa (FMT grant BE062). A T32 training grant to D.A. from the NIH (GM008666). Funding for collection of fibroblast cell lines was provided by a grant (IT1-06611) to J.G.G from CIRM. We are grateful to all of the families at the participating Simons Simplex Collection (SSC) sites, as well as the principal investigators (A. Beaudet, R. Bernier, J. Constantino, E. Cook, E. Fombonne, D. Geschwind, R. Goin-Kochel, E. Hanson, D. Grice, A. Klin, D. Ledbetter, C. Lord, C. Martin, D. Martin, R. Maxim, J. Miles, O. Ousley, K. Pelphrey, B. Peterson, J. Piggot, C. Saulnier, M. State, W. Stone, J. Sutcliffe, C. Walsh, Z. Warren, E. Wijsman). We appreciate obtaining access to phenotypic data on SFARI Base. Approved researchers can obtain the SSC population dataset described in this study (https://sfari.org/resources/autism-cohorts/simons-simplex-collection) by applying at https://base.sfari.org. The data reported in this paper are tabulated in the Supplementary Materials and archived at the National Database for Autism Research (DOI:10.15154/1340302), including the structural variant callset, our SV calling pipeline for use on AWS; forestSV, SV^2^and CNVplot software; raw sequence (FASTQ), alignment (BAM) and variant call (VCF) files from the REACH WGS dataset.

## Supplementary Tables

**Table S1.** Sample Information for 3,169 genomes

**Table S2.** Descriptive statistics of SV callset

**Table S3.** False Discovery rate of copy number variants across size ranges and filters

**Table S4.** SV callset permutations in functional elements

**Table S5.** Enrichment of known autism genes across pLI bins

**Table S6.** Group-wise Transmission/Disequilibrium Test analysis

**Table S7.** Variants in genes with pLI scores ≥90^th^ percentile

**Table S8.** Social responsiveness scale scores for parents, and children from the SSC cohort stratified on whether they carry SVs in variant-intolerant genes

**Table S9.** Fibroblast cell line expression analysis of *LEO1* and *MAPK6*

**Table S10.** De novo SVs

**Table S11.** Complex Mutation Clusters

**Table S12.** Case / control burden of de novo SVs

**Table S13.** Mosaic SV validation

**Table S14.** SVs in regions known to be associated with ASD

**Figure S1.**
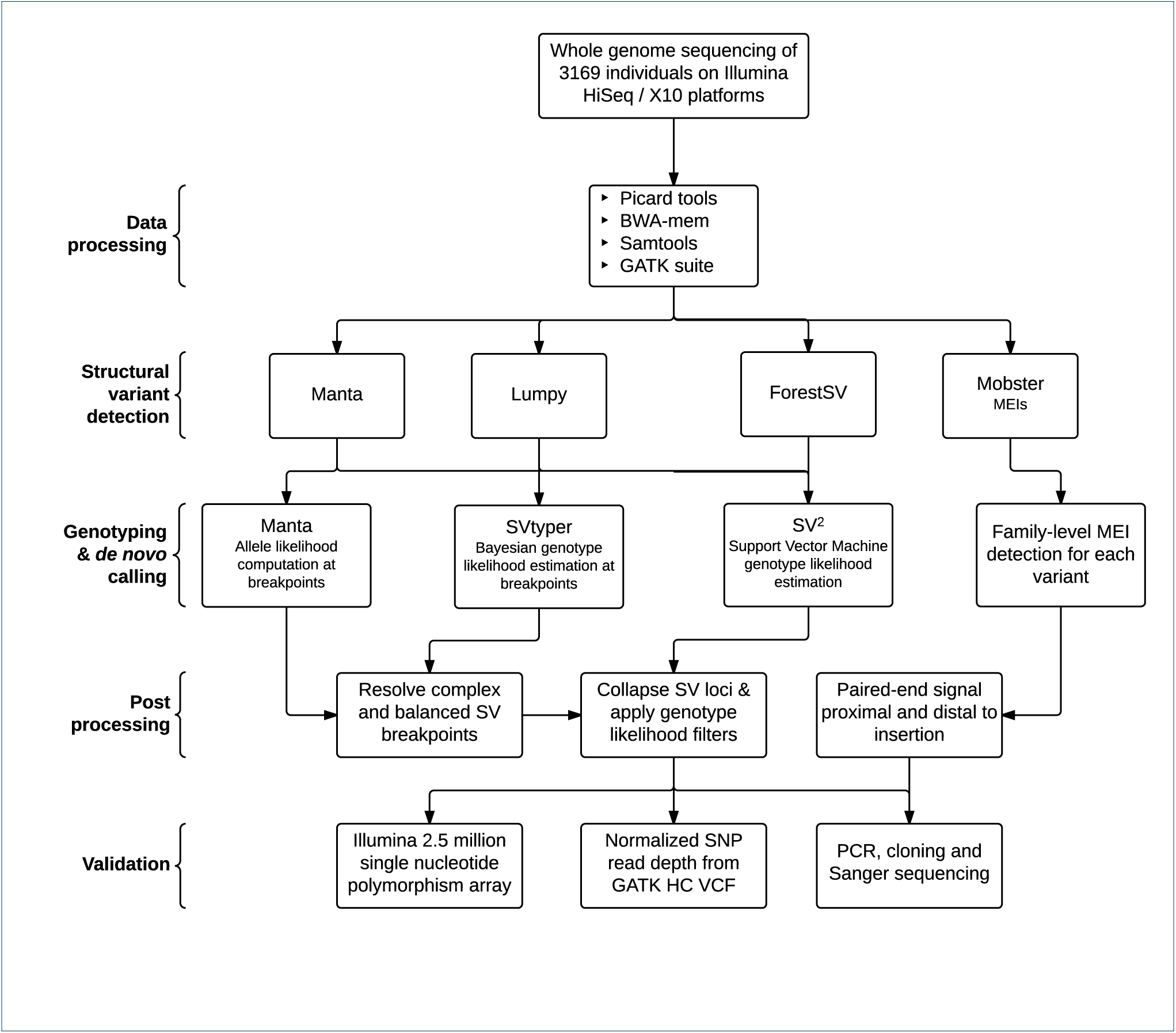
Structural Variant Discovery Pipeline. Flowchart detailing our custom pipeline for the discovery, genotyping, and validation of structural variants and de novo mutations. SV = Structural Variant; MEI = Mobile Element Insertion; PCR = Polymerase Chain Reaction.

**Figure S2.**
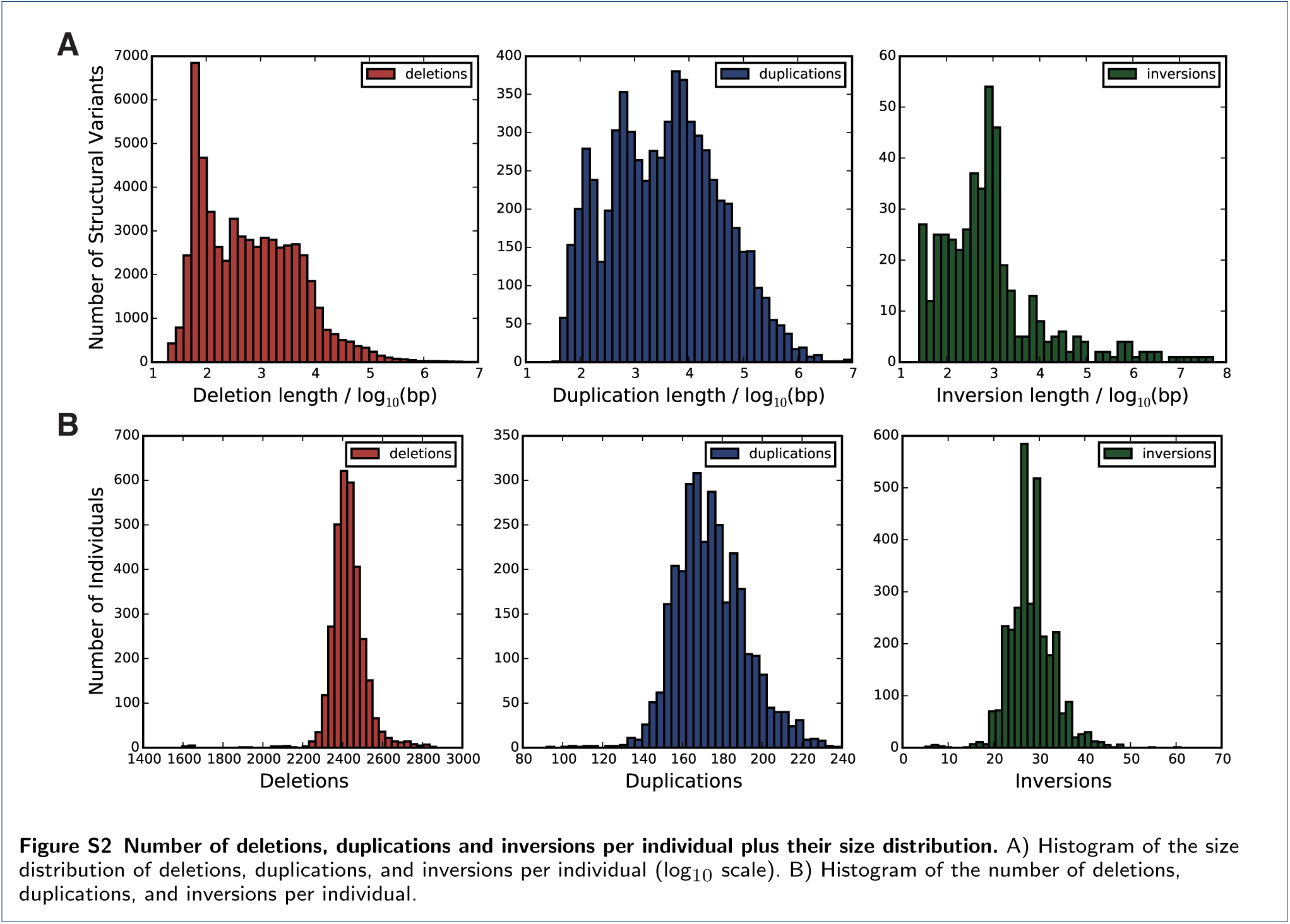
Number of deletions, duplications and inversions per individual plus their size distribution. A) Histogram of the size distribution of deletions, duplications, and inversions per individual (log_10_ scale). B) Histogram of the number of deletions, duplications, and inversions per individual.

**Figure S3.**
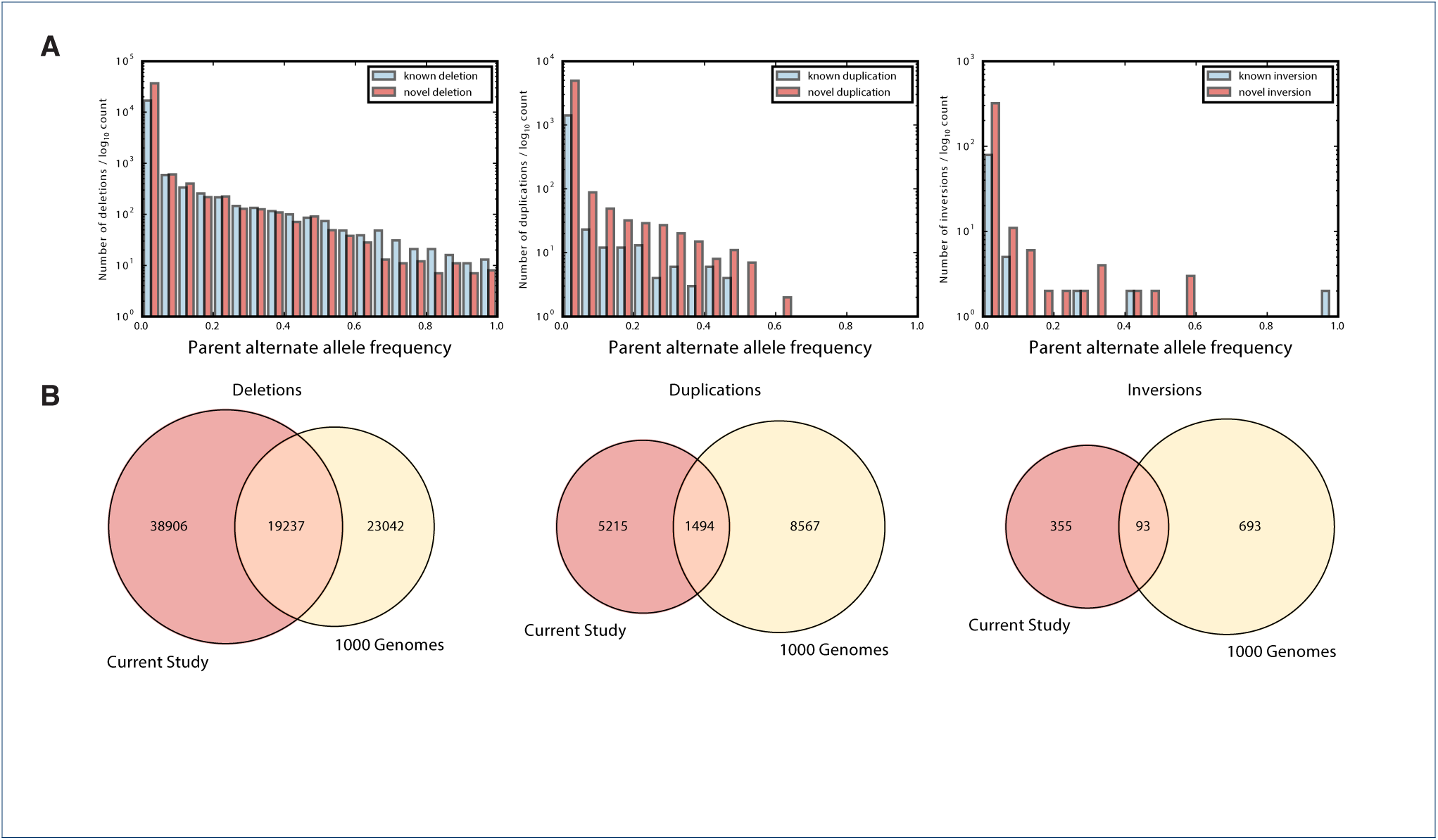
Callset overlap with 1000 Genomes Phase 3. A) Frequency of deletions, duplications, and inversions across parent allele frequency bins, stratified on known variants (from 1000 Genomes), and novel variants (detected only in this study). B) Venn diagrams of overlap of deletions, duplications, and inversions from our cohort with the 1000 Genomes

**Figure S4.**
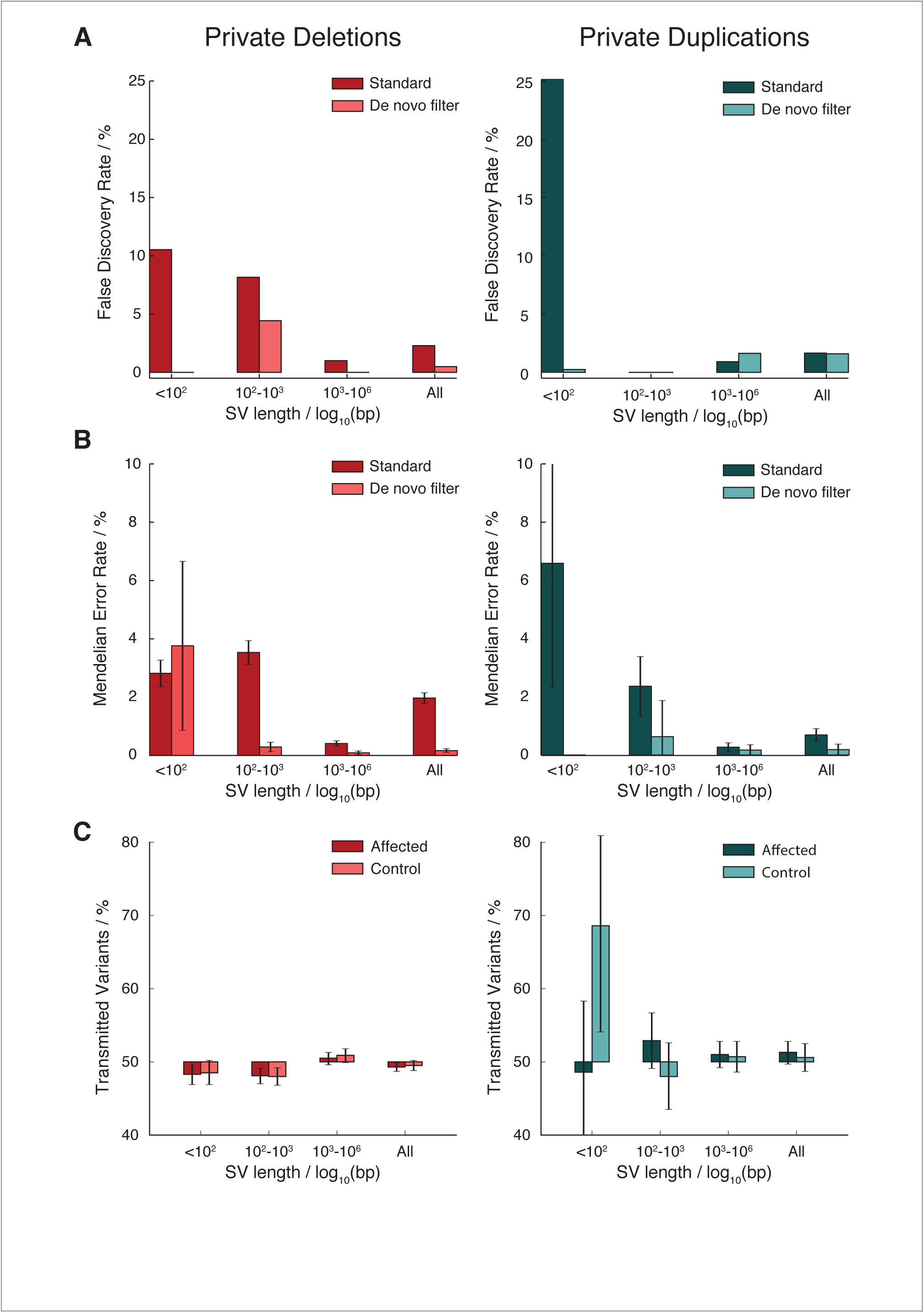
SV calling accuracy. Bar charts illustrating the A) FDR, B) Mendelian error rates, and C) variant transmission rates stratified on SV type (deletion and duplication) and SV length bins for private variants. Quality metrics are reported for all private SVs in the callset filtered based on SV^2^genotype likelihood at two levels of stringency (’standard’ and ‘de novo’). Whiskers represent 95% confidence intervals.

**Figure S5.**
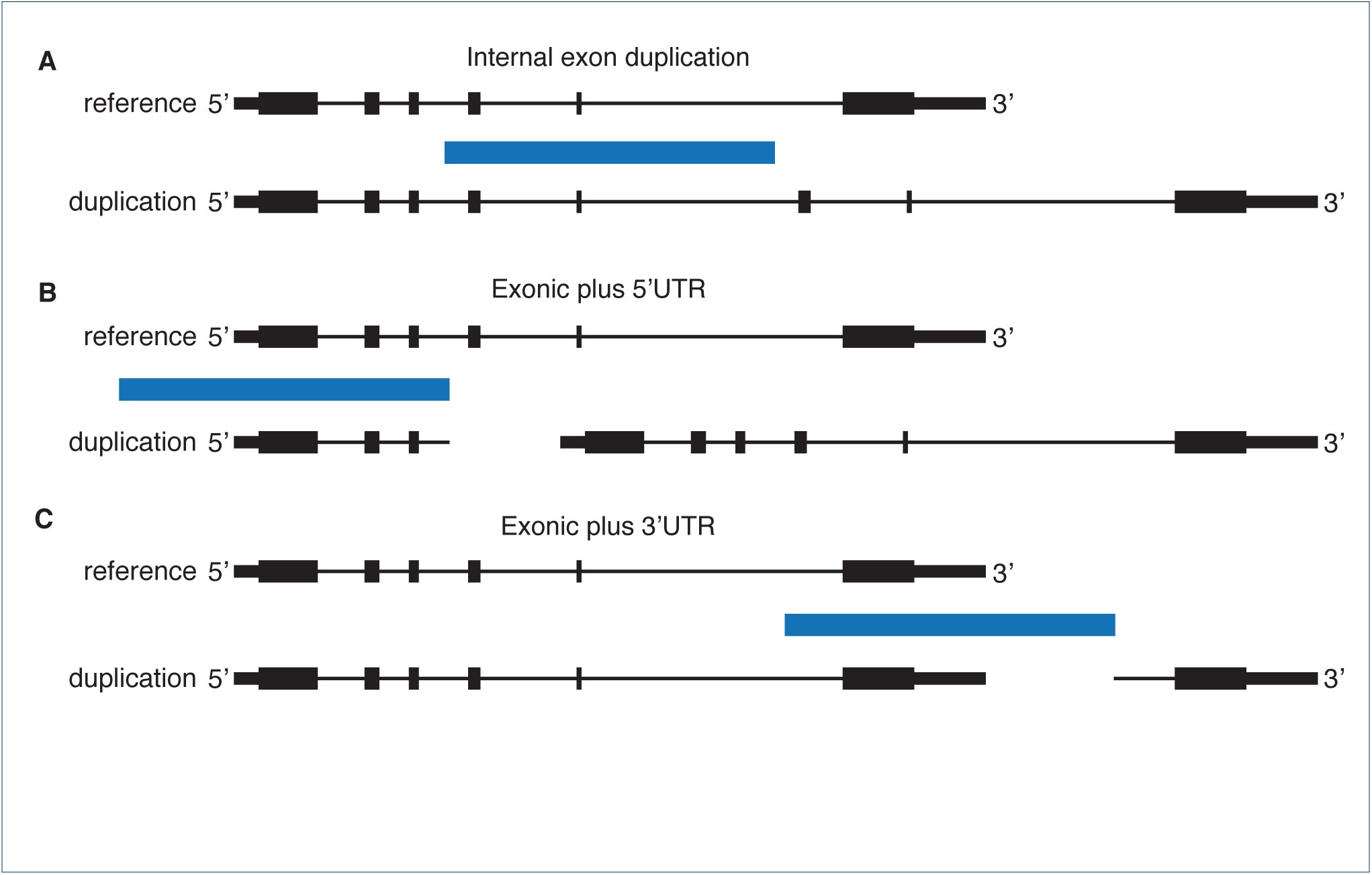
Functional impact of different classes of genic duplication. Diagrams illustrating how the functional impact of tandem duplications depends on their location within a gene, in each case the position of the duplication is shown by a blue bar, horizontal lines indicate intronic sequence, thin bars indicate UTRs and thick bars are protein coding exons; A) internal exon duplication, B) exonic duplication including the 5’UTR (and promoter), C) exonic duplication including the 3’UTR.

**Figure S6.**
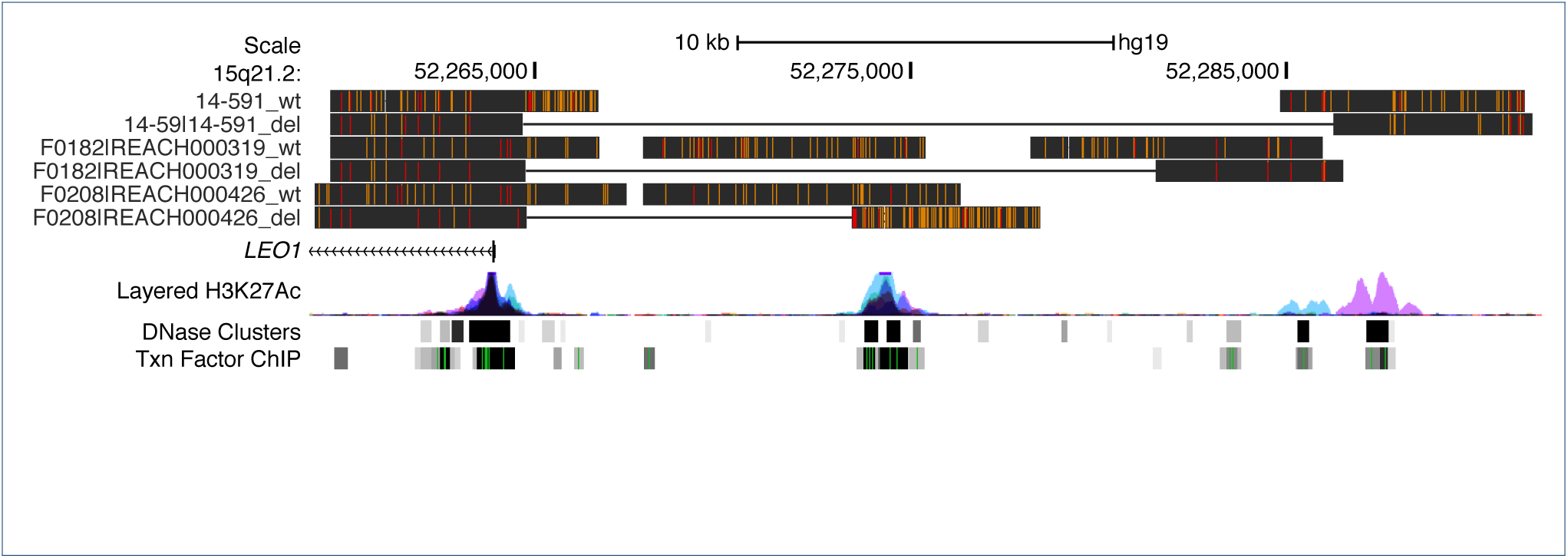
BLAT alignments from Oxford Nanopore sequencing of LEO1 deletions. UCSC genome browser image showing BLAT alignments of Oxford Nanopore long read sequences for three heterozygote deletions with corresponding wild type sequences. Black bars show alignments with yellow lines indicating indels and red lines SNPs. Wild type (wt) consensus contigs are shown within the breakpoint of the deletion. Deletion (del) contigs mapping either side of the breakpoints are linked with horizontal lines.

**Figure S7.**
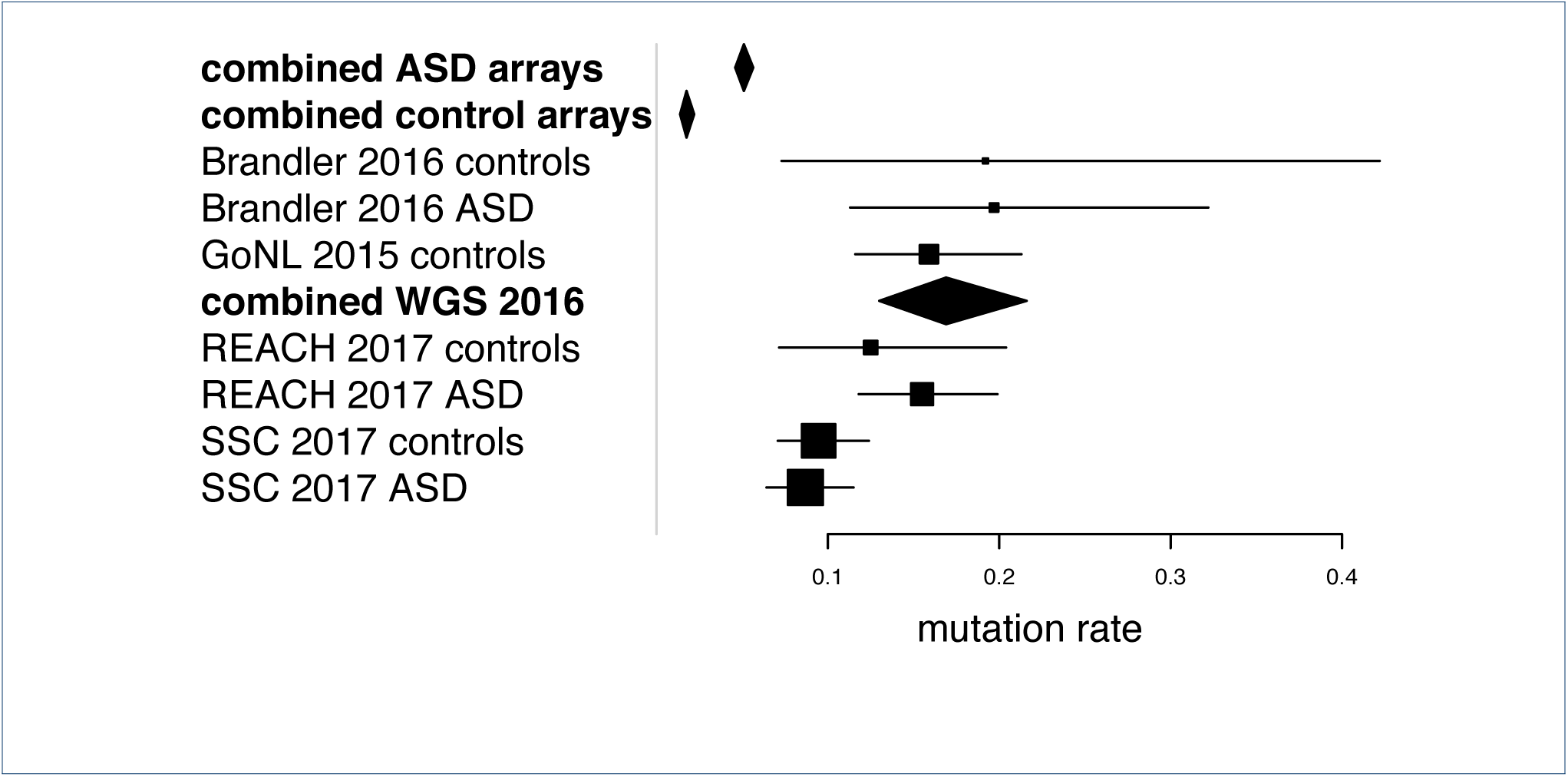
De novo mutation rate in the cohorts. Forest plot of the *de novo* mutation rate in the two cohorts from the present study (REACH 2017 and SSC 2017) compared to previous whole genome sequencing and microarray studies.

**Figure S8.**
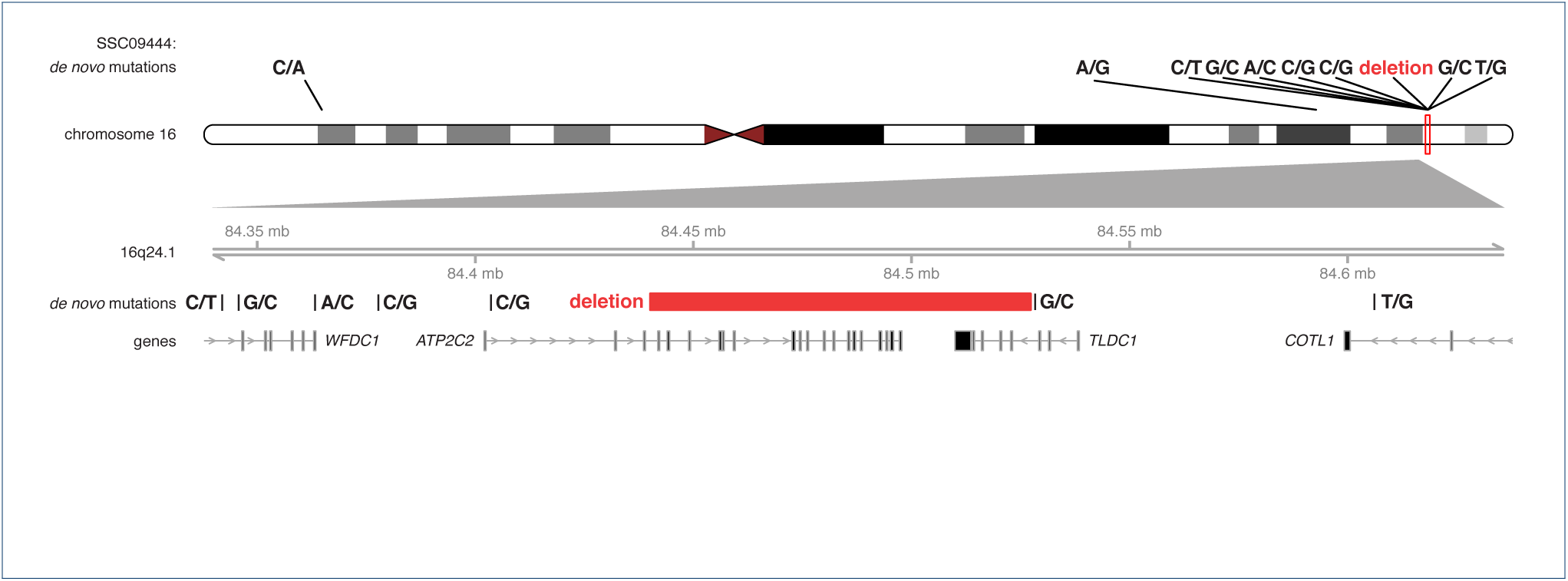
Mutational Clustering of SVs, Indels, and SNVs. One example of a complex mutation cluster are shown in the control individual from the SSC, SSC09444 (alternate ID: 13874.s1). The 300kb zoomed in locus below the ideogram shows the positions of de novo mutations relative to each other, an 82.3kb deletion is clustered with six SNVs upstream and two downstream of it. Gene tracks below the mutation show the longest transcript of each gene within the locus, with arrows indicating the strand and bars indicating the exons of genes.

**Figure S9.**
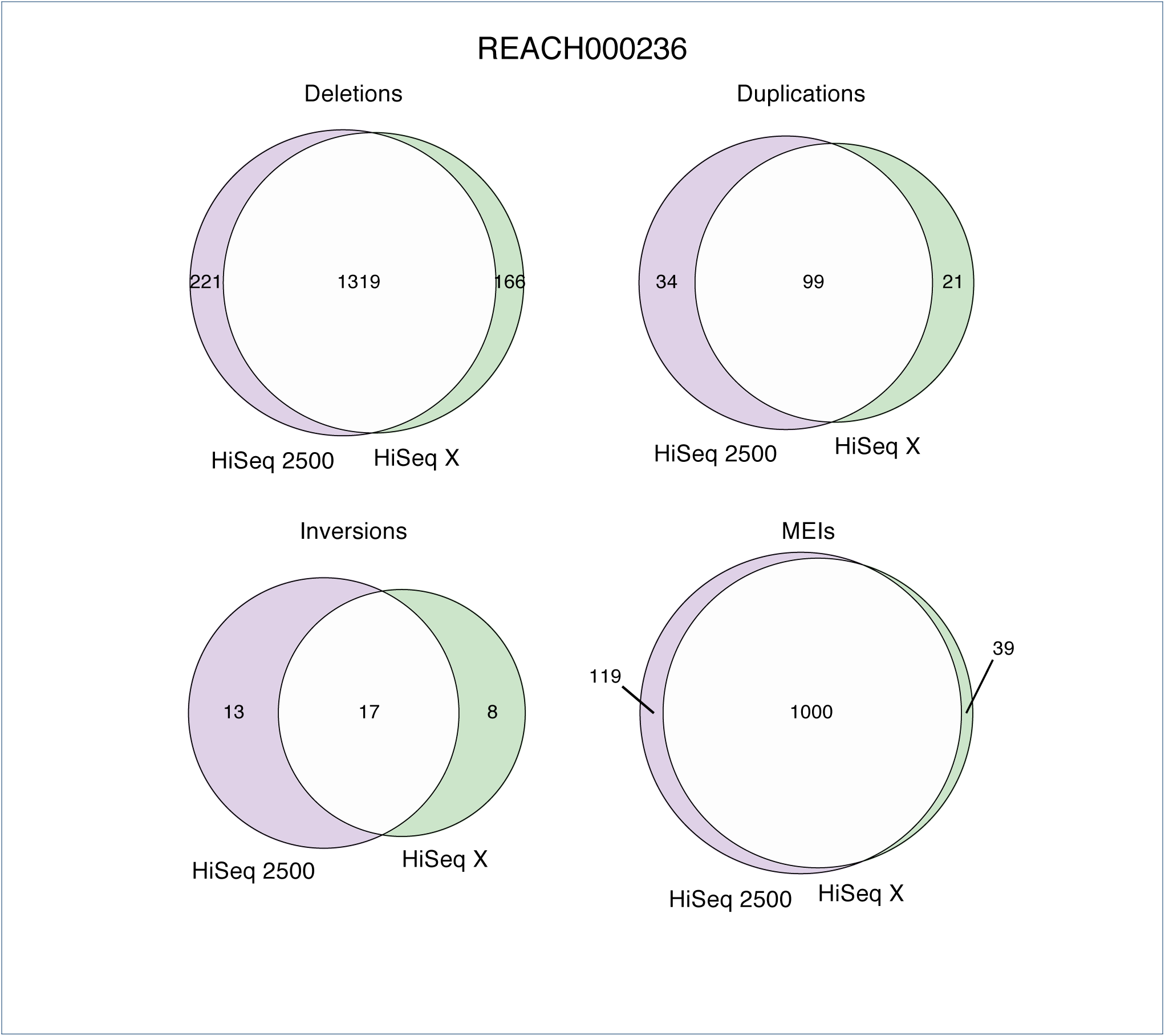
Overlap between SV calls made from one sample sequenced on two platforms. Sample REACH000236 was sequenced at 43X coverage on both the Illumina HiSeq 2500 with 100bp reads and on the Illumina HiSeq X with 150bp reads. Venn diagrams highlight the overlap for each SV type.

